# Whole genome functional characterization of RE1 silencers using a modified massively parallel reporter assay

**DOI:** 10.1101/2022.02.11.479757

**Authors:** Kousuke Mouri, Hannah B Dewey, Rodrigo Castro, Daniel Berenzy, Susan Kales, Ryan Tewhey

## Abstract

Both upregulation and downregulation by *cis*-regulatory elements help establish precise gene expression. Our understanding of how elements repress transcriptional activity is far more limited than activating elements. To address this gap, we characterized RE1, a group of transcriptional silencers bound by REST, on a genome-wide scale using an modified massively parallel reporter assay (MPRAduo). MPRAduo empirically defined a minimal binding strength of REST required for silencing (REST m-value), above which multiple cofactors colocalize and act to directly silence transcription. We identified 1,500 human variants that alter RE1 silencing and found their effect sizes are predictable when they overlap with REST binding sites above the m-value. In addition, we demonstrate that non-canonical REST binding motifs exhibit silencer function only if they precisely align two half sites with specific spacer length. Our results show mechanistic insights into RE1 silencer which allows us to predict its activity and effect of variants on RE1, providing a paradigm for performing genome-wide functional characterization of transcription factors binding sites.

## Introduction

Both inducing and repressing transcription by *cis*-regulatory elements (CREs) are crucial for the spatiotemporal responses controlling cell identity and function ^1^. More than a half century after the discovery of a repressive element acting on the *lac* operon ^2^, the rapid development of approaches to characterize CREs has revealed a multi-layered epigenetic landscape, highlighting a dynamic network of gene regulation responsible for multicellular control and environmental response ^3,4^. Detailed maps of histone modifications and chromatin accessibility have allowed us to annotate more than 800,000 candidate elements in the human genome that upregulate gene expression (i.e. enhancers and promoters) ^5^, while catalogues of elements that downregulate gene expression (i.e. silencers) are substantially smaller ^6^. Thus, comprehensive studies to understand the function of repressive elements are critical to understanding the full gene regulatory landscape ^7–9^.

Though most repressive elements remain poorly characterized, repressor element 1/neuron-restrictive silencer element (RE1/NRSE) is a well-defined group of silencers ^10,11^. RE1 is bound by Repressor element 1-silencing transcription factor (REST), also known as neuron-restrictive silencer factor (NRSF), which is a zinc finger transcriptional factor (TF) conserved through chordates ^12,13^. Although RE1 was initially discovered as a silencer for neuron-specific genes in non-neuronal cells, REST has also been found to have crucial roles in the brain and in the repression of non-neuronal genes ^14,15^. REST recruits Histone deacetylase (HDAC) complexes and Histone methyltransferase EHMT2/G9A to RE1 for silencing ^16–18^, mediated by cofactors including RCOR1/CoREST and SIN3A^16,19–21^. Localization of these cofactors has been demonstrated to vary among RE1s, suggesting different characteristics of RE1 dependent on the localization of cofactors ^22^. Our knowledge of TF interactions at RE1 can aid the understanding of silencer mechanisms but requires the systematic measurement of RE1 activities, which has not been done yet.

Massively parallel reporter assays (MPRA) are a high-throughput functional genomic platform designed to directly measure the activity of millions of CREs and can identify genetic variants that modulate their regulatory activity ^23–26^. To date, the majority of MPRAs have focused on identifying elements that enhance gene transcription, including enhancers and promoters, while repressive elements have remained understudied. This is partially due to the limited types of promoters used by most studies which are typically low activity, making it difficult to detect repression. While it has been demonstrated that using stronger promoters in MPRA helps to detect repressive elements, its sensitivity and specificity has not been evaluated systematically ^8,27^. In addition, it has been demonstrated that interactions between CREs can exhibit specificity depending on context including the cell-type and TFs involved ^28,29^. Thus, an appropriate selection of transcription-activating CREs is essential for testing repressive elements at-scale in reporter assays for the purpose of understanding their basic mechanisms and impact on disease. We sought to optimize MPRA to detect silencer activity at a scale.

## Results

### MPRAduo

To optimize the ability of MPRA to characterize silencer activity through the pairing of silencers with appropriate CREs, we developed MPRAduo which tests two CREs located in *cis* on the same reporter vector (Fig. 1A, Supplemental Fig. 1A). To associate tested elements and transcribed mRNA, we added a 10- and 20-nucleotide barcodes to the activating CRE (E) and silencer (S) modules respectively, then cloned them into two standard MPRA plasmid vectors using unique linker sequences (pΔGFP). As with standard MPRA libraries, we inserted a GFP ORF with a minimal promoter between the test sequence and barcode (we label the single libraries E and S). Next, we amplified the oligo-GFP-barcode cassette and cloned them into the reciprocal pΔGFP library resulting in combined E and S libraries for both alignments (duo libraries: SE and ES, according to the alignment of two elements from 5’ to 3’ in the upstream of a minimal promoter). Following transfection of libraries into cultured cells, we isolated the GFP mRNA and performed sequencing of the two barcodes in the 3’UTR to recover the elements and their orientation for downstream analysis to quantify the expression level (Methods).

**Fig. 1.**
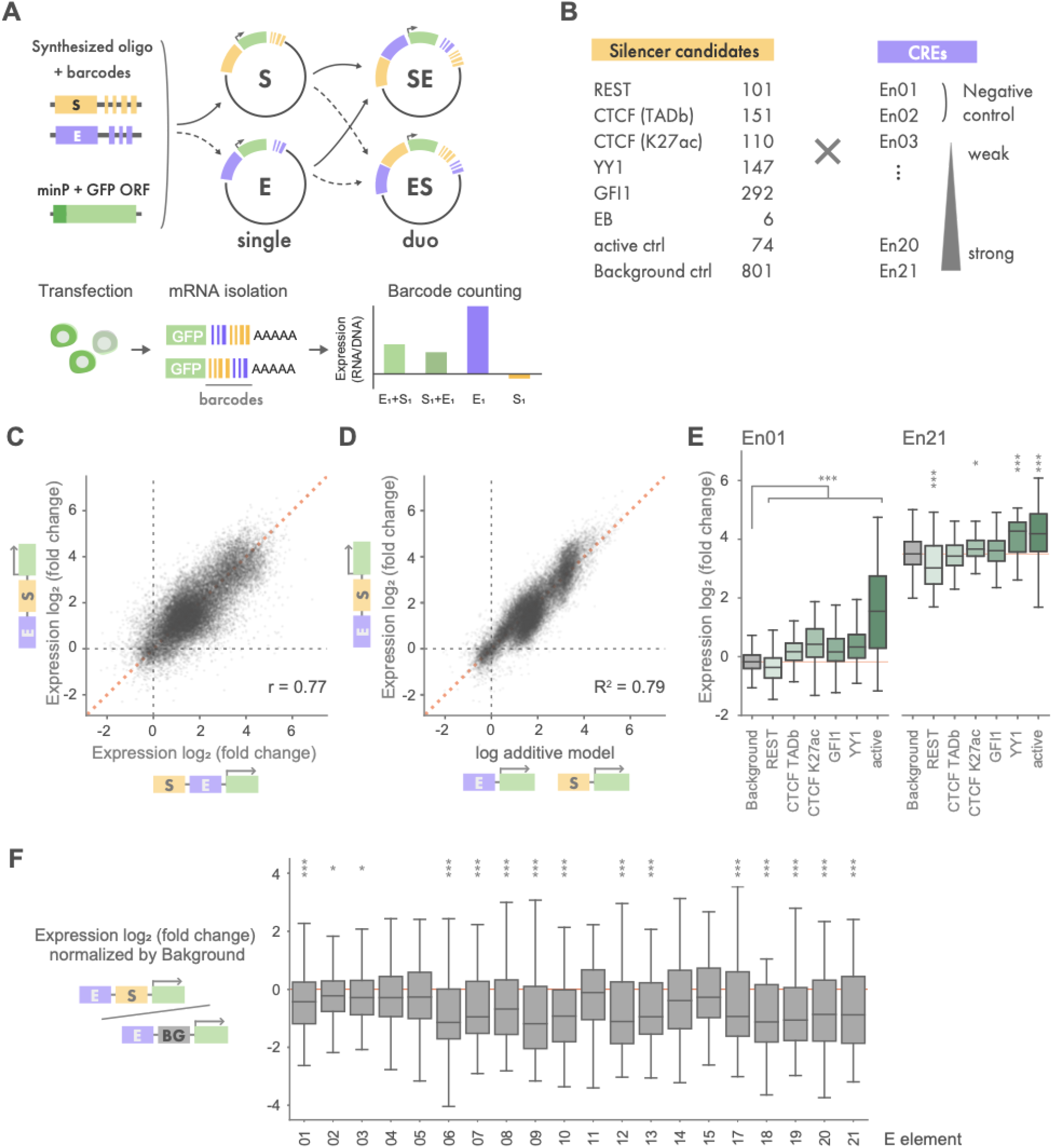
MPRAduo benchmarking. (A) Workflow of MPRAduo. Oligos are synthesized, barcoded, and cloned as single libraries with GFP ORF and a minimal promoter (minP). Then, the oligo and barcode sequences are amplified and cloned into another library (duo libraries). Libraries are transfected into cultured cell lines followed by isolation of GFP mRNA and sequencing. The aggregated count of mRNA reads are normalized by plasmid DNA counts. (B) Contents of the benchmarking library of MPRAduo. E elements are named in ascending order of their activity in MPRA of GM12878: En01 has the lowest expression while En21 has the highest. (C) Correlation of normalized expression levels (log_2_ of the mRNA/plasmid DNA ratio) between duo libraries. r indicates Pearson’s correlation. (D) Correlation between log additive model of single libraries and their observed duo values in SE library. (E) MPRA activity of each category of candidate silencers. Red line indicates the median of the negative control. (F) Activity of the combinations of RE1 and E elements in ES library normalized by the distribution of the corresponding combinations of RE1 and negative control. *: adjp<0.05, **: adjp<0.01, ***: adjp<0.001 by Mann-Whitney U-test (U-test) compared with background controls corrected using Benjamini Hochberg procedure (BH).

### MPRAduo Benchmarking

We used MPRAduo to identify CREs that, when tested in combination with putative silencers, respond to repressive effects. We selected 19 150-bp CREs previously tested by MPRA for evaluation in library E (E element) ^26^. E elements were chosen to represent a range of activity levels and TF binding. In addition, 2 negative controls with no reporter activity were included for a total of 21 unique sequences (Methods). For library S, 807 candidate silencer elements were selected from REST, CTCF (originating from topologically associating domain (TAD) boundaries or enhancer-like loci marked by H3K27ac), YY1, and GFI1 binding sites in the human genome, as well as the chicken HS4 sequence which is a well-known enhancer blocker (EB) and 5 human CTCF binding sites from previously validated EBs ^30,31^. In addition to the silencer candidates, we included 74 active controls and 806 matched background control sequences selected at random from the genome. Libraries E and S were constructed and both alignments were combined resulting in libraries ES and SE which contained in total 72,562 unique constructs.

We created two pools containing both single libraries and one duo library, and transfected each into GM12878 cells. Elements that had an activating effect in the single libraries showed high correlation between the two pools, indicating the system is highly reproducible across experiments (r = 0.77 for CREs, r = 0.65 for active control, Supplemental Fig. 1B). Duo libraries showed similar agreement between alignments (r = 0.77) and with the log additive model of the single library activity measurements (Fig, 1C and 1D, Supplemental Fig. 1).

MPRAduo showed significant repression by CTCF and RE1 (Fig. 1E, Supplemental Table. 5). CTCF binding sites at TAD boundaries significantly repressed the basal expression level of 4 E elements in the ES library and 9 E elements in the SE library (Supplemental Fig, 2A). RE1 showed the most significant repression in MPRAduo with 15 E elements in ES library and 13 E elements in SE library (Fig. 1F, Supplemental Fig. 2B). Overall, RE1 repressed activity of 12 E elements in both alignments with repression strength correlating with the basal level expression of E elements, although a few E elements, notably En11 and En15, were non-responsive to RE1 despite their medium to strong basal activities. These results demonstrate that MPRAduo can detect repression by RE1 and CTCF binding sites, and identifies CREs that improves the signal-to-noise ratio of silencers within a reporter assay.

### Whole genome RE1 screening

Encouraged by the ability of MPRAduo to characterize RE1 repression, we sought to comprehensively understand the mechanisms of RE1 activity genome-wide. We selected 8,436 RE1 sites containing a canonical REST binding motif and overlapping with a REST ChIP-seq peak in at least one of four human cell types: GM12878, K562, HepG2 and SK-N-SH (Fig. 2A). We also included 4,430 genomic sequences which overlap with a REST ChIP-seq peak in one or all of the four cell types but do not contain a canonical REST binding motif. To avoid evaluating promoters which are likely to increase expression in the reporter assay, we excluded all loci within 5-kb upstream of a transcription start site. Genomic sequences 200 bp in length containing the REST binding motif in its center (from 91st to 111th nucleotides) were synthesized and assembled in an ES library alongside 5 E elements tested from the benchmark set. We selected E elements to cover a range of activity levels, based on our benchmarking results, including one negative control (En02), one RE1 non-responsive CRE (En11), and 3 CREs which demonstrated significant repression by RE1 (En09, En19 and En21). We confirmed all four CREs are active, as marked by the presence of H3K27ac in the four cell types used in our screen ^32^.

**Fig. 2.**
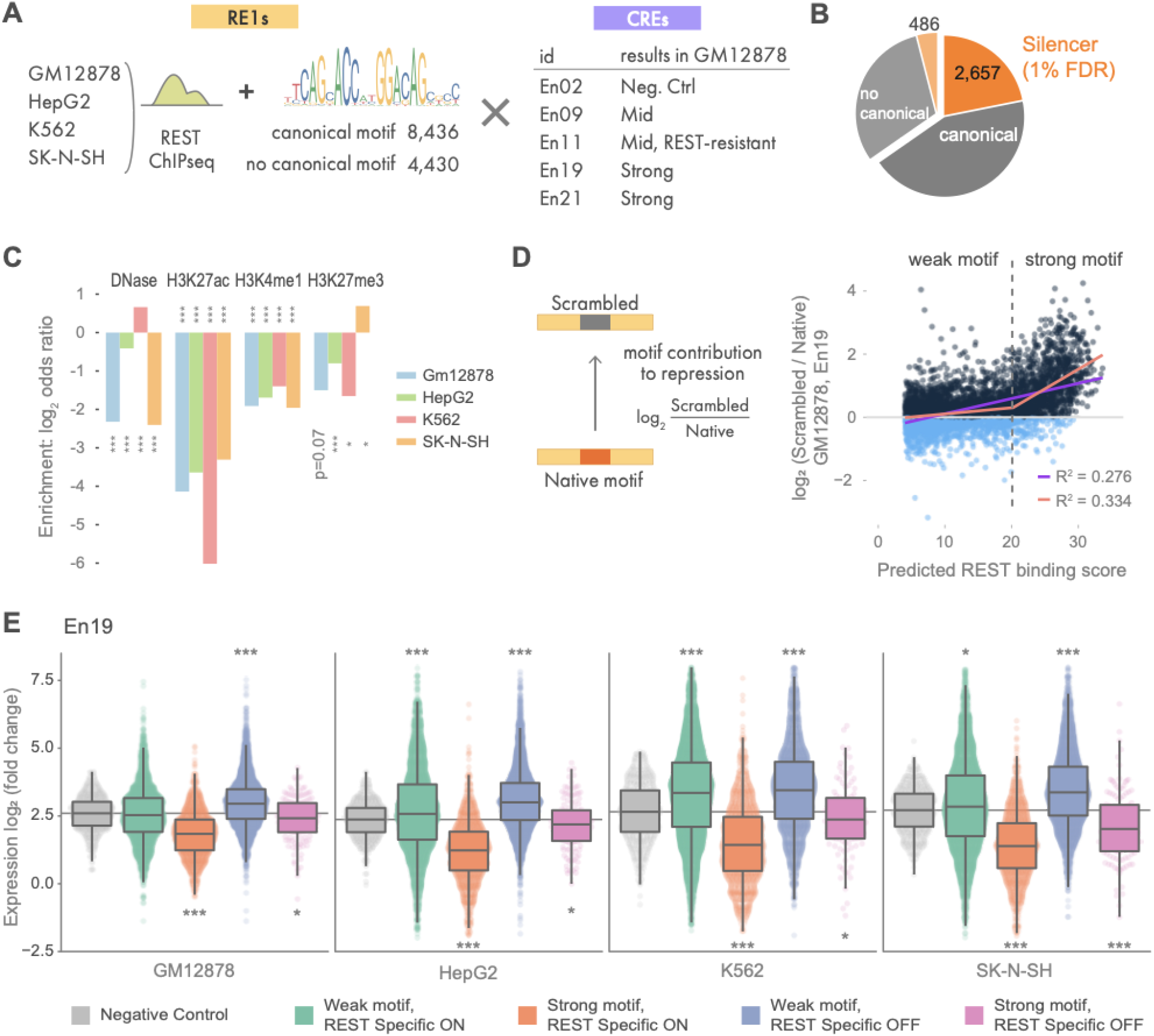
Whole genome RE1 screening. (A) Contents of the whole genome RE1 library. ChIP-seq peaks from 4 cell types with or without a canonical REST binding motif were combined with 5 enhancers. The names of enhancers correspond to the benchmarking set shown in Fig. 1. The motif shown is from JASPAR (MA0138.2). (B) Proportion of RE1 which showed repression with 1% of false discovery rate (FDR) in at least one cell/enhancer combination. (C) Log_2_(odds ratio) for enrichment of epigenetic marks in the strong silencers shown in Fig. 2B. **: p<0.01, ***: p<0.001 by Fisher’s exact test. (D) Correlation between predicted binding score of REST and motif contribution (log_2_(Scrambled / Native)) with En19 in GM12878. The purple line indicates linear regression and the orange line indicates piecewise linear regression. The dashed line indicates the change point determined by the piecewise linear regression. MPRAduo activity with En19 for each cell type. Specific-ON indicates RE1s bound by REST in each cell type and Specific-OFF indicates RE1s not bound by REST in the observing cell type but bound in other cell type(s). Grey line indicates the median of the negative control. *: adjp<0.05, **: adjp<0.01, ***: adjp<0.001 by U-test compared with negative controls corrected using BH.

We tested the whole-genome RE1 ES library in GM12878, HepG2, K562 and SK-N-SH cells (Supplemental Fig. 3). After correcting for the background activity of each E element we observed better correlations of the silencing activity of RE1s with canonical motif for each E element across cell types than across E elements in a single cell type suggesting generally REST activity within MPRAduo is more dependent on the genomic context than the cell type (Supplemental Fig. 3C). In total, 2,657 REST binding sites with a canonical motif (31%) and 486 without a canonical motif (10%) showed strong repression at a 1% FDR in at least one cell type and with one E element (Fig 2B). These 3,143 sequences were de-enriched of epigenetic marks for enhancers (H3K27ac & H3K4me1) in all four cell types tested with MPRAduo (Fig. 2C). We did not observe consensus enrichment for H3K27me3, a marker of Polycomb, in the 4 cell types.

To measure the precise contribution the 21 bp REST binding motifs have on the repression by each 200 bp RE1 sequence, we removed the REST motif for 5,866 RE1 sequence by scrambling the motif. An effect score for each REST motif was calculated by taking the difference of expression between the scrambled and native sequence (motif contribution) and compared with the predicted binding score of native sequence scored by FIMO ^33^ (Fig. 2D). While weak REST motifs did not contribute to repression, stronger motifs showed a clear contribution when paired with all E elements except for En02 and En11 in GM12878, the two elements which did not respond to RE1 in the pilot result. However, strong motifs contributed to the repression by RE1 when combined with the two E elements in the other three cell types, indicating the non-responsiveness of the two E elements for RE1 are cell-type specific. To determine the boundary between weak and strong motifs, we modeled the correlation between binding score and repressive activity using piecewise linear regression with a change-point estimation. The 18 of 20 E element-cell combinations with a clear shift of the correlation had an average change-point of 20.86 (ranged from 19.0 to 21.9) above which the slope of the regression dramatically increased (Supplemental Fig. 4, Supplemental Table 10); we used this average change-point as the boundary to delineate weak and strong REST binding motifs. The estimated values of the change point were close to each other (SD = 0.805), indicating that the change point is independent of cell types. Expanding the view to the whole 200 bp of the silencer elements, the RE1 sequences that have strong motifs and are bound by REST in the tested cell (Specific-ON) showed significant repression, while elements with weak motifs did not, and even showed a strong increase in expression for some sequences (Fig. 2E, Supplemental Fig. 6). These results demonstrate a clear boundary of the REST binding motif score (REST m-value) which determines RE1 silencer function in MPRAduo.

### Non-canonical binding motif requires precise arrangement and spacing of half sites

We next focused on exploring the 10% of the REST binding sites without canonical motifs that show repression by MPRAduo. The canonical 21 bp binding motif of REST consists of two half sites spaced 2 bp apart. Previous work has shown non-canonical motifs containing both half sites of the canonical motif with different combinations, orientations and spacer lengths are found in REST ChIP-seq peaks ^34,35^ (Fig. 3A). However, it is not clear how these arrangements of the half sites affect the silencer activity. To assess this, we identified half sites in the 4,430 REST binding sites without a canonical motif and annotated non-canonical pairs of half sites with summary binding score above the REST m-value determined by canonical motif. Our library includes 204 sequences containing an atypically spaced motif, 57 flipped, 52 convergent, and 54 divergent motifs as well as 509 sequences with a single half site. Atypically spaced motifs were the only configuration to show significant repression in MPRAduo (Fig 3B). Next, we compared the repression in MPRAduo of the different spacer lengths of atypically spaced motifs and found sequences with 8 and 9 bp spacers repressed expression while other distances showed no repressive effect (Fig. 3C and 3D). These results provide direct functional support that non-canonical REST binding motif requires precise alignment of two half sites with a specific spacer length of 8 or 9 bp for repressive activity.

**Fig.3.**
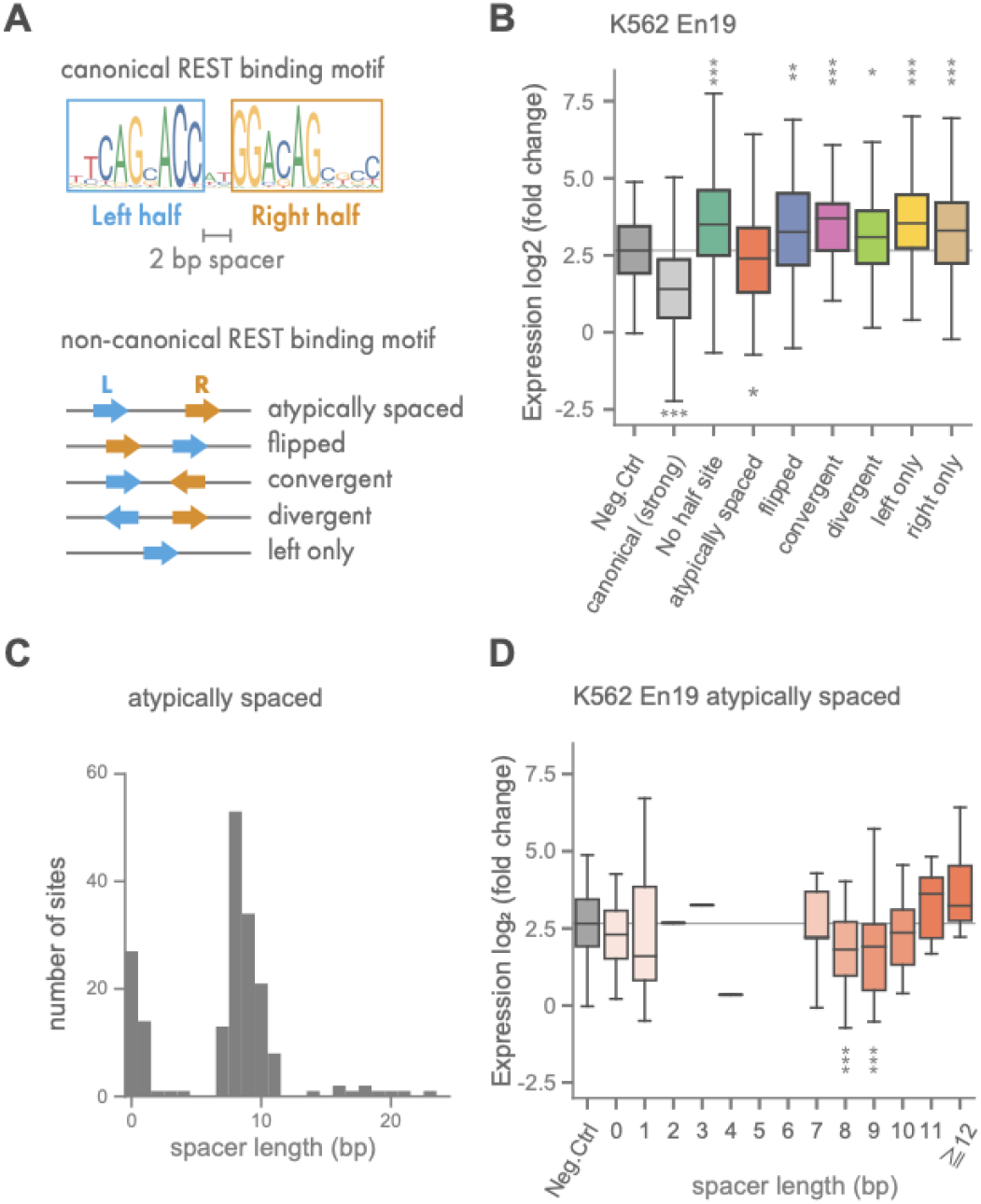
Spacer length of non-canonical REST binding motif impacts repression. (A) Structure of the canonical REST binding motif and classification of non-canonical motifs. Left and right half sites are separated by a 2-bp spacer in the canonical motif. Other motifs are classified according to the orientations and alignment of the two half sites. (B) MPRAduo activity of REST binding sites with or without canonical motif in K562 with En19. Grey line indicates the median of the negative control. (C) Distribution of spacer length of atypically spaced non-canonical motifs in the tested library. (D) Effect of spacer length of the atypically spaced non-canonical motif on expression level. Grey line indicates the median of the negative control. **: adjp<0.01, ***: adjp<0.001 by U-test compared with negative controls corrected using BH.

### Group of TFs colocalize with REST to facilitate silencer function

To classify additional co-factors of RE1, we sought to find TFs which may operate at RE1 in addition to REST. Using TF ChIP-seq data, we identified 329 TFs that are colocalized at our RE1 sequences with canonical REST motifs in K562 cells. For each TF, we separated RE1 sequences into four groups based on the binding of TF and REST as determined by ChIP-seq (e.g. TF+/REST+, TF+/REST-, TF-/REST+, & TF-/REST-). We calculated the difference of the median expression levels between these groups as measured by MPRAduo in K562 cells (Δmedian) (Fig, 4A). Using the direction of Δmedians between TF-positive and TF-negative groups with or without REST, we confidently placed 283 of 329 TFs into two of the four categories: (i) 269 TFs were associated with positive expression activity regardless of their colocalization with REST, and (ii) 14 TFs were associated with repressive activity when colocalized with REST but associated with positive activity when not colocalized with REST (Fig. 4B, red dots in the first and second quadrant respectively). No TFs were placed into category (iii), TFs associating with repressive activity regardless of REST colocalization, or category (iv), TFs associating with positive expression activity with REST and repressive activity without REST. Category (ii) includes AFF1, CHAMP1, CREB3, HINFP, MIER1, NCOA6, PTTG1, SREBF1, TEAD2, TRIP13, ZNF197, ZNF644 and ZNF766 as well as EHMT2, a known cofactor of REST, while category (i) includes two known cofactors: RCOR1 and HDAC2. All 14 TFs in category (ii) were significantly enriched in the RE1s with strong REST motifs compared to weak motifs while 191 of the category (i) TFs were significantly de-enriched, indicating that active RE1 recruits dual role repressor/activators and excludes REST-independent activators (Fig. 4C). We counted the number of category (ii) TFs localized at each RE1 sequence and observed a correlation with repressive activity that was dependent on REST, suggesting their recruitment is important for repression (Fig. 4D and 4E).

**Fig.4.**
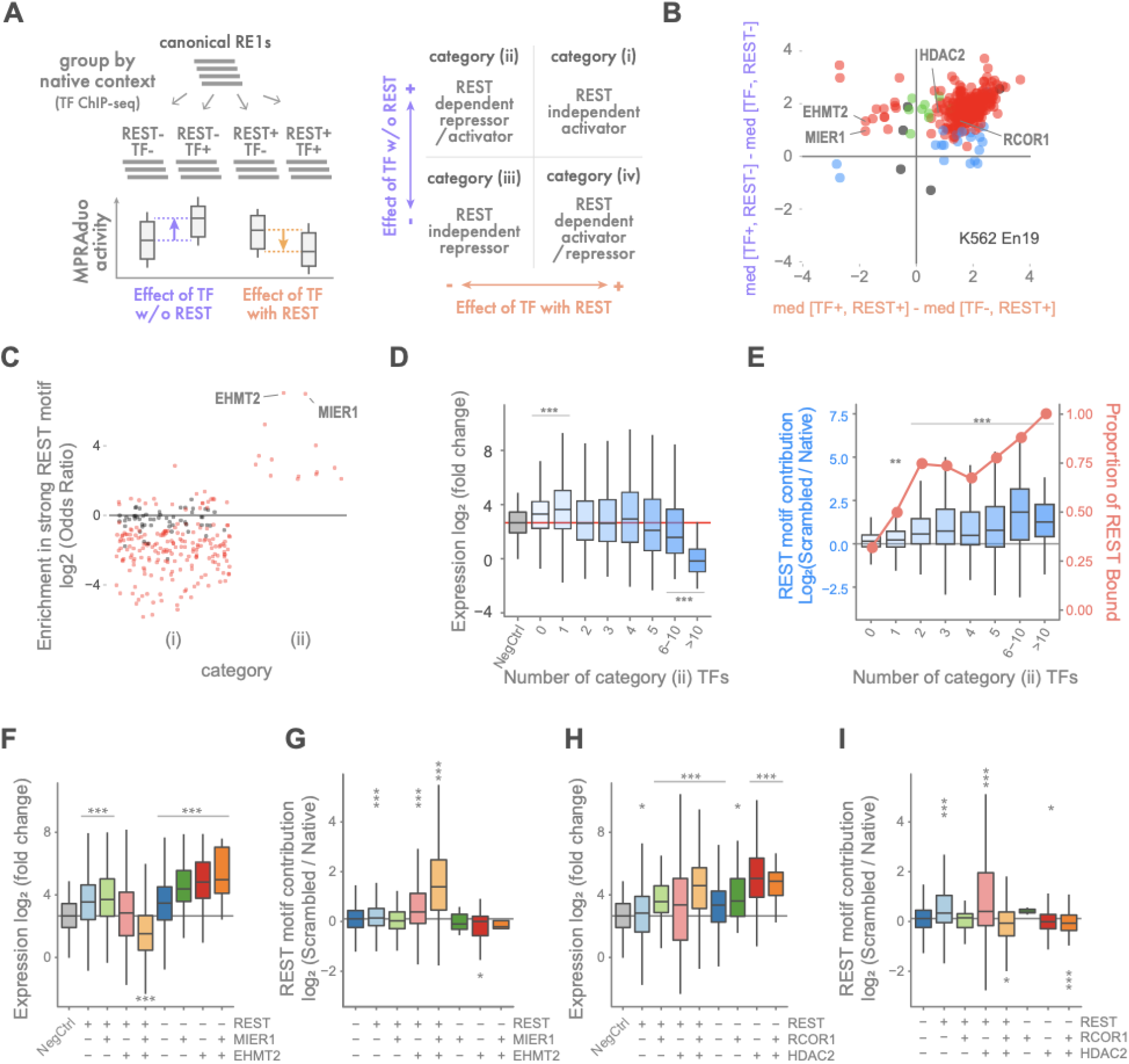
Screening and evaluation of REST cofactors. (A) RE1s with canonical motifs are grouped according to ChIP-seq of each TF and REST. Then differences of the medians of expression level between groups are plotted to categorize TFs into four which correspond to four quadrants of the plot. (B) Categorizing plot using TF ChIP-seq in K562 and reporter activity in K562 with En19. Red dots are TFs with significant differences despite REST colocalization. Blue and green dots are TFs with significant differences only with or without REST respectively (p<0.05 by U-test). (C) Enrichment of TF localization in K562 at RE1 with strong motifs compared to all tested canonical RE1. Red dots are significantly enriched TFs (FDR < 0.05 by Fisher’s exact test). (D) Correlation between number of localized category (ii) TFs and expression level of MPRAduo with En19 in K562. (E) Correlation between number of localized category (ii) TF and REST motif contribution with En19 in K562 (box plot, left axis). Red line indicates the proportion of REST binding sites of each bin (right axis). (F - I) Binding property of TFs and its effect on expression level (F and H) and REST motif contribution (g and i) with En19 in K562. *: p<0.05, **: p<0.01, ***: p<0.001 by U-test compared with negative controls (D, F, and H), RE1s bound by one TF (E), or triple negative group (G and I) corrected using BH.

We next focused on the 2 category (ii) TFs with the highest enrichment with strong REST motifs in our RE1 sequences; EHMT2 is recruited at RE1 by REST to suppress gene expression ^17^, and MIER1 interacts with EHMT2 and HDACs through its ELM2 and SANT domain ^21^. To evaluate the effect of MIER1 and EHMT2 colocalization on silencer function, we compared reporter activity and localizations of MIER1, EHMT2 and/or REST in the genomic context. RE1s associated with REST, MIER1, and EHMT2 showed significant repression in MPRAduo while the RE1s associated with REST and either MIER1 or EHMT2 showed no silencing effect (Fig. 4F). Furthermore, RE1s associated with MIER1 and/or EHMT2 but not REST increased reporter expression. The REST motif contribution measured by comparing scrambled and native motifs is highest in the RE1s associated with all three TFs, suggesting that MIER1 and EHMT2 require a REST motif to facilitate repression by RE1 (Fig. 4G). We recapitulated the correlation between repression and colocalization of MIER2 (a paralog of MIER1), EHMT2 and REST in HepG2 cells, which did not contain MIER1 ChIP-seq data, suggesting the redundant function of MIER proteins on RE1 (Supplemental Fig. 8).

Notably, we did not observe significant repression by RE1 sequences associated with HDAC2 and RCOR1 which were identified as category (i) TFs (Fig. 4H). However, REST motifs associated with REST and HDAC2 demonstrated a significant motif contribution, indicating that HDAC2 localized at functional RE1 silencers in the genome but it is not a sufficient marker for silencers (fig. 4I). Indeed, RE1s having strong REST motifs significantly reduced the expression level regardless of the association with RCOR1 and HDAC2, but when localized with the two cofactors showed stronger repression (Supplemental Fig. 9A). To confirm the difference of the cofactors localization in the native genomic context, we compared the ChIP-seq peaks in K562 and found that the weak motifs showed less occupancy by EHMT2 and MIER1 but a broad occupancy by RCOR1 and HDAC2, while all four cofactors showed higher occupancies at RE1s with strong motifs (Supplemental Fig. 9B). We also checked the gene expression of RE1 targets (defined as the nearest neighbor) and found the genes targeted by an RE1 associated with EHMT2 and REST showed significantly lower expression than the genes targeted by an RE1 without REST regardless of the motif strength while the repression by RE1 associated with HDAC2 and REST was shown only with strong motifs (Supplemental Fig. 9C). Overall, these results indicate that MIERs, EHMT2, and presumably other category (ii) TFs have a crucial role for the repression by RE1.

### RE1-modulating variants in Human genome

TF binding motifs enrich for fine-mapped variants associated with human disease ^36,37^. In order to understand the effect size and distribution of variants around REST binding motifs, we tested 1,450 variants of various allele frequencies located in the REST motif using MPRAduo as well as 2,348 variants located within 25 bp from the REST motif as control (Fig. 5A). We compared the expression level between alleles (“allelic skew”) and identified variants that showed significant differential expression between alleles (‘‘expression-modulating variants, emVar”). 642 emVars inside REST motif and 858 outside REST motif were detected and most of them were identified from K562 and SK-N-SH (FDR Δ 0.01, Supplemental Fig. 11, Supplemental Table 12). emVars were enriched inside of the strong binding motif compared to outside (odds ratio 2.89, p = 5.32 × 10^−12^ by Fisher’s exact test) but not enriched inside of the weak binding motif (odds ratio = 1.12, p = 0.171). In addition, variants falling within strong motifs showed greater allelic skew compared to weak motifs or variants falling outside a motif (Fig. 5B, Supplemental Fig. 12A). Allelic skew as measured by MPRAduo agreed with orthogonal measures of allelic activity, showing a strong correlation with the difference of the predicted binding score (delta binding score) between alleles at strong motif sites while those at weak motifs did not correlate (Fig. 5C, Supplemental Fig. 12B). These results demonstrate the importance of first identifying variants that fall into strong motifs with binding score above the REST m-value prior to considering the effect of the variant on REST binding.

**Fig. 5.**
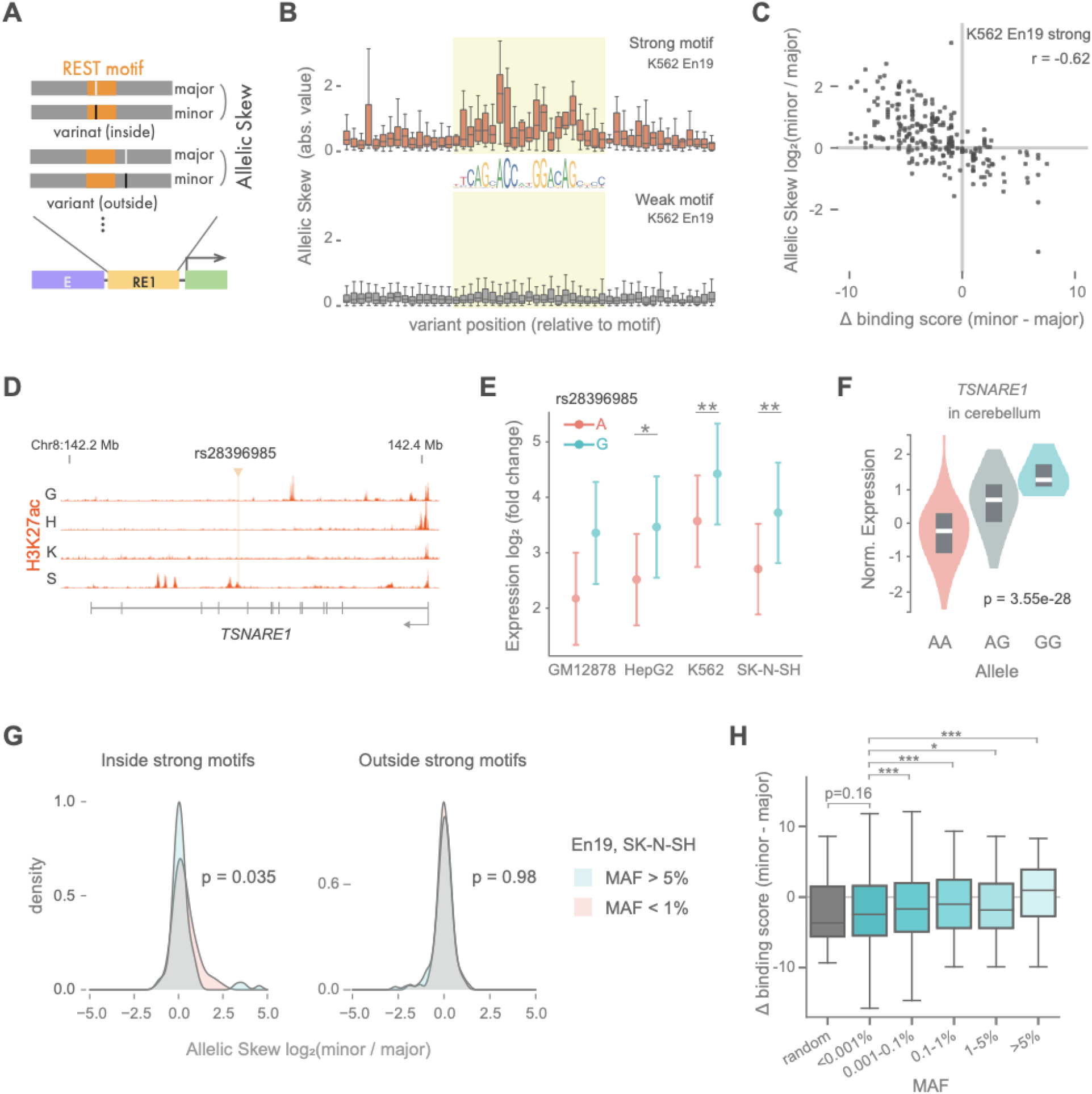
Strong motif enriches strong variants. (A) Allelec skew to test the effect of the variants inside or outside the canonical REST binding motif. Both reference (Ref) and alternate (Alt) alleles are tested in MPRAduo and the difference of the activities is measured as allelic skew. (B) Distribution of absolute value of allelic skew along variant position relative to REST binding motif. Allelic skew with En19 in K562 for variants around strong motif (top) and weak motif (bottom) are plotted. Light green highlights the canonical REST binding motif (JASPAR MA0138.2). (C) Correlation of allelic skew and delta binding score. Allelic skew with En19 in K562 is plotted against the difference of predicted binding score between alleles. (D) Position of rs28396985 and H3K27ac ChIP-seq signal in *TSNARE1* locus. G: GM12878, H: HepG2, K: K562, S: SK-N-SH The y-axis is from 0 to 32 for all tracks. (E) MPRAduo activity of RE1 containing each allele for rs28396985 with En19. Error bars indicate standard error. *: < 1% FDR (F) eQTL signal of rs28396985 for *TSNARE1* expression in human cerebrum. Normalized expression level of *TSNARE1* is plotted for each allele. (G) Distribution of allelic skew with En19 in SK-N-SH in different MAF. (H) Correlation between delta binding score and MAF of variants at genome-wide strong REST binding motif. *: p<0.05, ***: p<0.001 by U-test compared to <0.001% MAF corrected using BH.

One example of an emVAR impacting a strong REST motif is rs28396985 which is in the intron of *TSNARE1* (Fig. 5D, Supplemental Fig. 12C). rs28396985 is located at the *cis*-regulatory element bound by CTCF without any active marks as measured by H3K27ac or H3K4me3 ^5^. The G allele showed a significant increase of the expression relative to the A allele (Fig. 5E) which is in agreement with the predicted REST delta-binding score (34.1 for the A-allele and 30.4 for the G-allele). eQTL results by GTEx showed that expression of *TSNARE1* is increased by the G allele in multiple tissues including cerebellum and spleen, indicating a significant effect of the variant *in vivo* that is in agreement with the effect measured using MPRAduo (Fig. 5F).

### RE1-modulating variants show allele frequency differences

To understand how genetic variants in REST binding sites impacted function, we interrogated the relationship between allele frequency and REST activity. For the majority of the major alleles, as estimated by global allele frequency, we observe a stronger predicted REST binding scores and repressive effects by MPRAduo than the corresponding minor allele (Fig. 5C). As we selected variants in REST ChIP-seq peaks from only four cell types, this effect may be due to ascertainment bias against rare or low frequency alleles that create a REST binding site. However, even after binning variants based on low (<1%) and moderate to high (>5%) allele frequency we still observed a noticeable effect that was strongest at low frequency (Fig. 5G). To perform an unbiased and exhaustive assessment, we next identified all known variants, both common and rare, which are predicted to create a REST binding motifs in the human genome regardless of their overlap with an existing REST ChIP-seq peak. We identified 10,069 variants where either allele contributes to a strong REST binding motif, and then compared the predicted binding score between the major and minor alleles. We used the delta binding score as a proxy for altered REST activity due to the strong correlations we observed with MPRAduo.

In agreement with the imbalance observed by MPRAduo, lower minor allele frequency (MAF) variants had significantly lower delta binding score (which corresponds to high allelic skew by MPRAduo) than higher MAF variants which were more evenly distributed between positive and negative values (Fig. 5H). To identify a baseline expectation, we created random *de novo* variants which exhibited negative delta binding scores similar to the lowest MAF group (< 0.001%). These findings suggest that low frequency variants represent a random process and alleles at increased allele frequency in the population may undergo selective pressures against the disruption of established REST binding sites.

## Discussion

In this study, we characterized whole genome RE1 silencers using MPRAduo which enables us to maximize our ability to detect repressive effects. We empirically identified a motif-intrinsic value (m-value) for REST: RE1 where a binding score above this m-value establishes an effective silencer likely through the recruitment of cofactors including MIERs and EHMT2. We note that weak binding motifs below the REST m-value overlap with REST ChIP-seq peaks. However, they colocalize more often with activators rather than cofactor candidates in the genome and more often increase reporter expression level. These observations suggest weak REST motifs are less likely to play a role in actively silencing gene expression and may only have weak effects ^41^. TFs in category (ii) are repressive when localized with REST but associated with an increase of expression when they appear without REST implying that some TFs are capable of both repressive and activating function which is dependent on their surrounding context ^42,43^. Such a dual role has been demonstrated for EHMT2 with the binding site and phosphorylation playing key roles ^44,45^. Our results suggest REST, when bound to a strong motif, may assist in switching the function of TFs to facilitate silencer activity.

Our results show that a spacer length of 8 or 9 bp is required for silencing activity in non-canonical REST, concordant with the enrichment of the non-canonical motif with those specific spacer length in REST ChIP-seq peaks ^34,35^. We did not find any consensus sequences in the 8 or 9 bp spacers, suggesting that the specific spacer length, and not the content, is important for the structural interaction between DNA and REST. Although the majority of TF binding motifs are described using a Positional Weight Matrix (PWM) with fixed length, some TF binding motifs with variable spacer length have been discovered ^46,47^. Our results emphasize the importance of motif discovery with variable gap length and functional validation.

We determined that accumulation of category (ii) TFs at RE1 in the genome correlates with repression measured by MPRAduo, suggesting that category (ii) TFs associate with REST at RE1 to facilitate silencer function. The MIER family has been demonstrated to physically interact with REST as well as EHMT2, suggesting that MIERs help the interaction between REST and EHMT2 in a protein complex ^48^. ZNF644 is a Zinc finger transcription factor which also binds to EHMT2, providing additional support for a role of category (ii) TFs as mediators at RE1 ^49^. Category (ii) also includes proteins not previously associated with the RE1 complex such as TRIP13, an AAA+ ATPase which plays a role during meiosis ^50^. We confirmed the localization of both ZNF644 and TRIP13 are associated with repression by RE1 in MPRAduo and repression of target gene expression in their genomic context (Supplemental Fig. 10). However, direct evidence of interactions between REST and category (ii) TFs within the genome are required for further verification.

Although genome-wide association studies are a powerful tool to find variants associated with traits, it is a challenge to isolate causal variants from other variants in tight linkage disequilibrium ^36,51^. Chromatin marks and functional assays, including MPRA, provide strong evidence for nominating causal variants however, intersecting variants only with TF binding motifs typically does not enrich for causal variants as effectively as those methods ^52^. This work demonstrates that empirically estimating an m-value for each TF as a stringency filter could play an important role for identifying sequences that have clear functional activity, and furthermore, to aid the prioritization of variants that impact human health.

## Methods

### Vector design

Two different vectors were designed to be used for single libraries: vectors A (pMPRAv3:Δluc:ΔxbaI, addgene #109035) and P (pMPRAduo:Δorf, addgene pending) contain unique cloning sites corresponding to different oligo adapter sequences (A:5’ACTGGCCGCTTGACG [150 or 200 bp oligo] CACTGCGGCTCCTGC3’, P: 5’ACTGGCCTCGCTTGC [150 or 200 bp oligo] CCCTGGCCGACCTGG3’) and have AsiSI or PmeI recognition sequences, respectively, inserted between the oligo and barcode to insert GFP and the other library. In the benchmarking library, vector A was used to clone library S with 1,687 sequences and vector P was used to clone library E with 21 sequences. In the whole genome RE1 library, vector P was used for library S with 24,000 sequences and vector A was used for library E with 5 sequences.

### Oligo synthesis and barcoding

Oligos for libraries S were synthesized by Agilent Technologies (benchmarking library) and Twist Bioscience (whole genome RE1 library) as 230 bp sequence containing 200 bp of unique sequence flanked by 15 bp adapter sequences. After synthesis, 20 bp barcodes and additional adapter sequence were added by 4 or 6 cycles of 12 PCR reactions each 50 μL in volume containing oligos, 25 μL of Q5 NEBNext Master Mix (NEB, M0541) and 0.5 μM forward and reverse primers (Integrated DNA Technologies, IDT) (primers 1 and 3 for the benchmarking library and 4 and 6 for the whole gnome RE1 library, Supplemental Table 1) cycled with the following conditions: 98°C for 30sec, 4 or 6 cycles of (98°C for 10 sec, 60°C for 15 sec, 65°C 45 sec), 72°C for 5 min. Chicken *HS4* sequence was amplified from pBluescriptII[attB/Ins1] (addgene #74100), adding 20 bp barcodes and adapter sequences by 6 cycles of PCR reactions 50 μL in volume in the same method as the benchmarking library.

Oligos for library E were synthesized as 180 bp sequences containing 150 bp of genomic context and 15 bp of adapter sequences (IDT) and cloned into the pMPRAduo:Δorf vector without barcodes, sequence verified, and individual clones were selected. Individual plasmids were linearized by PmeI (NEB, R0560), 180 bp sequence were amplified to add 10 bp barcodes and additional adapter sequences using a 6 cycles of PCR reaction in 20 μL volume containing 10 μL of Q5 NEBNext Master Mix and 0.5 μM forward and reverse primers (IDT) (primers 4 and 5, Supplemental Table 1) cycled with the following conditions: 98°C for 30sec, 6 cycles of (98°C for 10 sec, 60°C for 15 sec, 65°C 45 sec), 72°C for 5 min. The PCR products were purified using 1× volume of AMpure XP (Beckman Coulter, A63881) and eluted with water. The purified amplicons were equalmolar pooled and assembled into vector P for benchmarking library or separately assembled into A-vector for Whole-genome REST library.

### Vector assembly of single libraries

To assemble the single Δorf library, 1 μg of PCR product containing barcoded oligos were inserted into 1 μg SfiI (NEB, R0123) digested empty vector A or P by gibson assembly NEBuilder HiFi DNA Assembly Master Mix (NEB, E2621) in an 80 μL reaction. After 1 hr of incubation at 50°C, DNA was purified using a Monarch PCR & DNA Clean up kit (NEB, T1030) and eluted in 12 μL of water.

Assembled vectors of library S and *cHS4* were mixed with a 1500:1 ratio of molarity. The mixture electroporated into 100 μL of 10-beta E.coli (NEB, C3020K, 2kV, 200 ohm, 25 μF) in order to achieve a transformation efficiency equal to 300-times the number of unique oligos sequences (target CFU: 500K). The electroporated bacteria was immediately split into ten 1 mL aliquots of outgrowth medium (NEB, B9035S) and incubated at 37°C for 1 hr then independently scaled up in 20 mL of LB supplemented with 100 μg/mL of carbenicillin (Teknova, C8001) and incubated on a shaker at 37°C for 9.5 hr; after outgrowth, the cultured bacteria was pooled prior plasmid purification (Qiagen, 12943 or 12963). Four of the aliquots were sampled and plated with serial dilutions after 1 hr recovery to estimate transformation efficiency.

Library E was transformed into 5-alpha E.coli (NEB, C2987H), recovered in SOC medium (NEB, B9020S) and incubated at 37°C for 1 hr then diluted and plated onto LB agar supplemented with 100 μg/mL of carbenicillin (Teknova, L1010) and incubated overnight at 37°C. Approximately 2100 colonies were harvested by washing plates with LB, followed by plasmid purification (Qiagen, 12943). To construct the whole genome RE1 binding library, individual clones for each of the 5 library E sequences were cultured in 20 μL of LB with 100 μg/mL of carbenicillin overnight at 37°C, equal volumes of the clones were combined together and the pool expanded in 20 mL of LB with 100 μg/mL carbenicillin for 9 hr followed by plasmid purification (Qiagen, 12943). Approximately 250 colonies in total were harvested.

To construct the mpra:gfp single library, 10 μg of Δorf library plasmid was digested with 100 units of AsiSI for vector A or PmeI for vector P (NEB, R0630 or R0560) at 37°C overnight. A GFP open reading frame (ORF) with a minimal promoter and partial 3’ UTR was amplified from pMPRAduo:minP:GFP (addgene pending) in 1600 μL volume containing 800 μL of Q5 High-Fidelity 2X Master Mix (NEB, M0492L) and 0.5 μM forward and reverse primers (primers 7 and 8 for vector A and primers 9 and 10 for vector P, Supplemental Table 1) cycled with the following conditions: 98°C for 30sec, 20 cycles of (98°C for 10 sec, 60°C for 15 sec, 72°C 45 sec), 72°C for 5 min. The PCR product was purified using 1.5× volume of AMpure XP and inserted by gibson assembly using 3 μg of linearized Δorf library and 9 μg of the GFP amplicon in 100 μL total volume for 90 minutes at 50°C. Assembled vectors were purified using 1× volume of AMpure XP and a secondary digest performed with 40 U of AsiSI or PmeI, 5 U of RecBCD (NEB, M0345), 10 μg BSA, 1 mM ATP, 1× NEB Buffer 4 at 37°C overnight followed by purification with a Monarch PCR & DNA Clean up kit using 12 μL of water for elution. 3 μL of mpra:gfp plasmid was transformed into 50 μL of 10-beta cell by electroporation (2 kV, 200 ohm 25 μF). The electroporated bacteria were recovered and cultured with 100 mL of LB supplemented with 100 μg/mL of carbenicillin in the same way of Δorf library.

### Vector assembly of duo libraries

To assemble the duo library for the benchmarking set, oligos and barcodes including GFP ORF with minimal promoter were amplified from the single mpra:gfp library of plasmid A or P by PCR using 50 μL reaction volumes containing 1 ng of plasmid, 25 μL of Q5 NEBNext Master Mix and 0.5 μM forward and reverse primers (IDT) (primers 11 and 12 to amplify vector A and 13 and 14 to amplify vector P, Supplemental Table 1) cycled with the following conditions: 98°C for 30sec, 12 cycles of (98°C for 10 sec, 60°C for 15 sec, 65°C 1 min), 72°C for 5 min. The amplified cassettes were inserted by gibson assembly using 4 μg of the amplicons and 2 μg of Δorf library plasmid, linearized by AsiSI or PmeI, in a 200 μL reaction incubated for 90 min at 50°C followed by AMpure XP purification using 75 μL of water for elution. Total eluted volume was digested by incubation with 50 U of AsiSI, 50 U of PmeI, 5 U of RecBCD, 10 μg BSA, 1 mM ATP, 1× NEB Buffer 4 at 37°C overnight and purified by Monarch PCR & DNA Clean up kit using 12 μL of water for elution.

For the whole genome RE1 library, 220 ng of barcoded oligos were directly inserted into 2 μg of AsiSI-digested single Δorf library using gibson assembly in a 200 μL reaction incubated for 60 min at 50°C and purified by Monarch PCR & DNA Clean up kit using 12 μL of water for elution. The purified ligation product was electroporated into 100 μL of 10-beta E.coli (NEB, C3020K, 2kV, 200 ohm, 25 μF) in order to achieve a transformation efficiency equal to 200-times the number of unique oligos combinations (target CFU: 4.8 M). The electroporated bacteria was immediately split into ten 1 mL aliquots of outgrowth medium and incubated at 37°C for 1 hr then independently scaled up in 20 mL of LB supplemented with 100 μg/mL of carbenicillin and incubated on a shaker at 37°C for 9.5 hr; after outgrowth, the cultured bacteria was pooled prior plasmid purification (Qiagen, 12963). 20 μg of Δorf duo library plasmid was digested with180 U of PmeI and 20 U of AsiSI at 37°C overnight. A GFP open reading frame (ORF) with a minimal promoter and partial 3’ UTR was amplified and inserted by gibson assembly using 1.6 μg of linearized Δorf duo library and 5.3 μg of the GFP amplicon in 250 μL total volume for 90 minutes at 50°C. Assembled vectors were purified using 1× volume of AMpure XP and a secondary digest performed with 40 U of AsiSI or PmeI, 5 U of RecBCD (NEB, M0345), 10 μg BSA, 1 mM ATP, 1× NEB Buffer 4 at 37°C overnight followed by purification with AMpureXP using 40 μL of water for elution. 6 μL of mpra:gfp plasmid was transformed into 200 μL of 10-beta cell by electroporation (2 kV, 200 ohm 25 μF). The electroporated bacteria were recovered and expanded in 3000 mL of TB (Teknova, T0315) supplemented with 100 μg/mL of carbenicillin for 16 hr at 30°C followed by plasmid purification (Qiagen, 12991).

### MPRA transfections

Lymphoblastoid cells (GM12878) were grown in RPMI 1640 (Thermo Fisher Scientific, 61870036) supplemented with 15% of FBS (Thermo Fisher Scientific, A3160402) maintaining a cell density of 2-10 × 10^5^ cells/mL. 10^7^ cells were mixed with 10 μg of plasmid and electroporated in a 100 μL volumes of RPMI with the Neon transfection system (Thermo Fisher Scientific, MPK5000) using 3 pulses of 1200 V for 20 msec. In total, 10 × 10^7^ cells were used for each replicate of the benchmark librarys and 50 × 10^7^ cells were used for each replicate of the whole genome RE1 library. HepG2 cells were grown in DMEM (Thermo Fisher Scientific, 10566024) supplemented with 10% of FBS maintaining a cell density of 2-5× 10^5^ cells/cm^2^. 10^7^ cells mixed with 5 μg of plasmid were electroporated in a 100 μL volumes of Resuspension Buffer R with the Neon transfection system using 1 pulse at 1200 V for 50 msec each. In total, 15 × 10^7^ cells were used for each replicate. K562 cells were grown in RPMI 1640 supplemented with 10% of FBS maintaining a cell density of 2-10 × 10^5^ cells/mL. 10^7^ cells mixed with 5 μg of plasmid were electroporated in a 100 μL volumes of Resuspension Buffer R with the Neon transfection system using 3 pulses of 1450 V for 10 msec. In total, 15 × 10^7^ cells were used for each replicate. SK-N-SH cells were grown in DMEM supplemented with 10% of FBS maintaining a cell density of 3-15 × 10^5^ cells/cm^2^. 10^7^ cells mixed with 10 μg of plasmid were electroporated in a 100 μL volumes of Resuspension Buffer R with the Neon transfection system using 3 pulses of 1200 V for 20 msec. In total, 15 × 10^7^ cells were used for each replicate. All cell lines were recovered from the culture medium 24 hr post-transfection by centrifugation, washed 3 times with PBS, and frozen at −80°C in Buffer RLT supplemented 40 mM of DTT.

### RNA extraction and cDNA synthesis

Total RNA of the transfected cells was extracted using RNeasy Maxi (Qiagen, 75162) following the manufacturer’s protocol including the on-column DNase treatment. Total RNA was secondarily digested by 20 U of Turbo DNase (Thermo Fisher Scientific, AM2238) in 1.65 mL of total volume for 1 hr at 37°C. The digestion was stopped by the addition of 15 μL of 10% SDS and 150 μL of 0.5 M EDTA, followed by incubation at 70°C for 5 min. To capture GFP mRNA,1200 μL of Formamide (Thermo Fisher, 4311320), 600 μL of 20× SSC (Thermo Fisher, 15557044) and 2 μL of Biotin-labeled GFP probe (primers 15 - 17, Supplemental Table 1) were added directly to the stopped DNase reaction mixture and incubated for 2.5 hr at 65°C with rotation. 400 μL of Dynabeads Streptavidin C1 (Thermo Fisher, 65002) was prewashed, eluted to 500 μL of 20× SSC and added to the reaction followed by agitation on a HulaMixer (Thermo Fisher, 15920D) at room temperature for 15 min. The magnetic beads were then washed once with 1× SSC and twice with 0.1× SSC and 50 μL of water was added along with 1 U of SUPERase In (Thermo Fisher, AM2694). Beads were treated with 2 U of Turbo DNase at 37°C overnight and the digestion was stopped with the addition of 1 μL of 10% SDS followed by purification with RNA clean XP purification beads (Bechman Coulter, A63987) using 37 μL of water for elution. cDNA was synthesized from the purified DNase-treated GFP mRNA using SuperScript III with a 1 μM final concentration of primer specific to the 3’ UTR of GFP (primer 18, Supplemental Table 1) at 47°C for 80 min. Synthesized cDNA was purified by AMpure XP and eluted in 30 μL of EB (Qiagen, 19086).

### Sequencing library construction and Illumina sequencing

To pair barcodes with oligo sequences Δorf plasmid was amplified by PCR in a total reaction volume of 200 μL containing 400 ng of plasmid DNA, 200 μL of Q5 NEBNext Master Mix and 0.5 μM of forward and reverse primers (primers 19 and 20 for plasmid A and 21 and 22 for plasmid P, Supplemental Table 1) cycled with the following conditions: 98°C for 30sec, 5 cycles of (98°C for 10 sec, 62°C for 15 sec, 72°C 30 sec), 72°C for 2 min. PCR products were purified using 1× volume of AMpure XP and eluted in 30 μL of EB. Illumina indices were added to each sample by amplifying 20 μL of the elution in a 100 μL of PCR reaction with 50 μL of Q5 NEBNext Master mix and 0.5 μM of forward and reverse primers (primers 25 and 26, Supplemental Table 1) cycled with the following conditions: 98°C for 30sec,6 cycles of (98°C for 10 sec, 62°C for 15 sec, 72°C 30 sec), 72°C for 2 min. Indexed samples were purified using 1× volume of AMpure XP, eluted in 30 μL of EB and sequenced using 2×250 bp chemistry on an Illumina MiSeq instrument at the Jackson Laboratory.

cDNA tag sequencing libraries were amplified by PCR each 100 μL in volume containing 10 μL of cDNA, 50 μL of Q5 NEBNext Master Mix and 0.5 μM of forward and reverse primers (primers 23 and 24 in Supplemental Table 1) cycled with the following conditions: 98°C for 30sec, 6-13 cycles of (98°C for 10 sec, 62°C for 15 sec, 72°C 30 sec), 72°C for 2 min. The cycle number was estimated by qPCR with 10 μL of the same reaction and 1:60000 diluted SYBR Green I (Thermo Fisher, S7563) and 1 μL of cDNA or 1 μL of diluted plasmid DNA used as a standard curve for the qPCR. To construct tag sequencing libraries of the plasmid pools, plasmids were diluted based on the qpcr results to mirror the cDNA samples, and amplified using the same PCR conditions and cycles as the cDNA. PCR products of the cDNA and plasmid libraries were purified using 1× volume of AMpure XP and eluted in 30 μL of EB. Illumina indices were added to each sample by amplifying 20 μL of the elution in a 100 μL of PCR reaction with 50 μL of Q5 NEBNext Master mix and 0.5 μM of forward and reverse primers (primers 25 and 26, Supplemental Table 1) cycled with the following conditions: 98°C for 30sec,6 cycles of (98°C for 10 sec, 62°C for 15 sec, 72°C 30 sec), 72°C for 2 min. Indexed samples were purified using 1× volume of AMpure XP, eluted in 30 μL of EB and sequenced using 2×150 bp chemistry on an Illumina NextSeq 550 or 1×150 bp S1 chemistry on an Illumina NovaSeq instrument at the Jackson Laboratory.

### Pre-Processing of reads

The first of the pre-processing steps works to create a map between the barcodes and oligos. Paired-end 250 bp reads from the sequencing of both single libraries were merged into single amplicons using Flash2 ^53^ (v2.2.00, flags: -M 200), then, the positions of the UMI, barcode, and oligo from each amplicon were identified by using the 3’ linker sequences of the barcode/oligo and the 5’ linker sequences of the UMI/barcode/oligo (Supplemental Fig. 13). The barcode-oligo pairs were then aligned to the original oligo sequences using minimap2 ^54^, and the resulting SAM output was filtered for mapping quality. Barcodes were sorted and parsed to remove barcodes mapping to multiple oligos and organized into a dictionary of barcode-oligo pairs.

The second pre-processing pipeline extracts counts from the replicated tag sequences. In the benchmark set, the tag sequences were based on paired-end 150-bp reads; reads were first merged into single amplicons using Flash2 for each replicate. In the whole genome RE1 set, the tag sequences were 150-bp single-end reads, so this step was unnecessary. In both sequencing sets the reads were sorted into single and duo libraries based on the number of bases between the tail of the GFP and 3’ end of the sequence. Here, regardless of whether or not there were single barcodes in the replicate sequences, any duo barcode sets that had less than 110 bp from the GFP tail to the 3’ end were classified as singles. The barcodes were extracted and the ones that were present in the dictionary, or in the case of duo barcodes in both dictionaries, were included in the count table.

### Full analysis

Datasets were filtered based on the number of barcodes observed for an oligo’s count (≧20 benchmark, ≧10 whole genome RE1) and the average number of DNA counts across replicates (≧100 benchmark, ≧20 whole genome RE1). Since the benchmark set included single libraries, and utilized two sequencing runs, the oligos were additionally filtered to ensure the single libraries contained the same oligos across all libraries, and that the duo oligos were only made up of oligos that were present in the filtered single oligo libraries. After filtering the oligos, a DESeq-based analysis ^55^ followed by a summit-shift normalization was performed. On the whole genome RE1 library, this was expanded with a cell-type specific analysis that was adjusted based on the summit shift normalization. To identify “expression-modulating variants” (emVars) in the whole genome RE1 library, the difference between the log2FoldChange of the alternate and reference allele for each replicate within a cell type were compared using the student’s t-test similar to previous approaches ^26^. In order to effectively compare between the two runs in the benchmark set, a quantile normalization ^56^ of the log2FoldChanges was performed across libraries.

For the benchmarking library, 69,456 constructs were recovered from all four libraries (95.7%), 55,077 passed the filters (75.9% of all constructs). After filtering, 28,297 and 25,455 constructs were included from SE and ES libraries respectively, with 25,222 combinations captured in both alignments and used in the downstream analysis (Supplemental Table 4). For the whole genome RE1 library, 115,830 (96.52% of all constructs) constructs on average were recovered across cell types. After filtering, 105,794 (88.2% of all constructs) constructs on average across cell types were used in the downstream analysis (Supplemental Table 6).

### Silencer Selection of Benchmark Library

To generate silencer candidates for the benchmark set, ChIP-seq peaks from ENCODE overlapping with TF binding motif were selected with a cut off binding score for REST (ENCFF048JKT, Factorbook score 5.95) and YY1 (ENCFF967ACD, Factorbook score 3.9). CTCF sites (ENCFF002DAJ, Factorbook score 4.95) were intersected by Factorbook motifs and ChIP-seq peaks of H3K27ac (ENCFF411MHX) or 40kb from TAD boundaries ^57^. Putative GFI1 binding sites were selected for inclusion based only on having a Factorbook score higher than 0.9. Selected binding sites were expanded to 200 bp by centering the TF motif in the test sequence and extending the genomic sequence on either end. TF binding sites which were located 5 kb from transcription start sites (TSS) annotated by Ensembl were removed. Fifty thousand random genomic sequences were selected by using bedtools (version 2.29.2). After removing sequences located 5 kb from TSS, 806 random sequences matching the distribution of GC content for the TF binding sequences were selected as random negative controls.

### E elements Selection

To select the 19 E elements for the benchmarking set, non-coding human elements which were previously tested for activity in GM12878 and HepG2 cells by MPRA were used ^26^. 602 elements which had significant activity (-log10Padj > 4) in both cells and also overlapped at least one transcription factor binding annotation in ENCODE for GM12878 or HepG2 were selected. These elements were separated into 24 clusters using k-means (Scikit learn, version 0.24.2). From 12 out of 24 clusters which had more than 20 elements, 19 E elements were picked up to have diverse expression levels in MPRA and diverse TF binding based on ChIP-seq peaks in GM12878 from ENCODE. Two negative controls which did not show expression in MPRA or do not have active marks (H3K27ac, H3K4me1, H3K4me3) were also selected from *Tewhey et al 2016*.

### Whole Genome RE1 library

To select sequences for the whole genome RE1 library, narrow peak data sets of ChIP-seq for REST in 4 cell types (GM12878: ENCFF048JKT, ENVFF677KJB, HepG2: ENCFF153JLK, ENCFF854KPC, K562: ENCFF895QLA, ENCFF558VPP, SK-N-SH: ENCFF781PAL, ENCFF946MYA from ENCODE) were combined together. A 21 bp REST binding motif (Factorbook from ENCODE) was intersected with the ChIP-seq peaks and expanded 90 bp to 5’ and 89 bp to 3’ by bedtools. Elements located within 5 kb upstream from a TSS were removed. To identify reference sequences without canonical motifs, 1000 cell specific ChIP-seq peaks for each cell type were randomly selected from the narrow peaks which did not intersect with the REST binding motif or reside within 5 kb upstream of a TSS. For GM12878, only 934 peaks fit this criteria and all were included in the library. In addition, all 496 sequences without canonical motifs which overlap ChIP-seq peaks in all 4 cell types were added to the library. These peaks were trimmed or expanded to 200 bp, keeping the center of the peak at the center of oligos. Alternate alleles of 2,865 human variants with 1% or higher MAF and randomly selected 1,000 human variants with less than 1% of MAF in at least one of three populations (eastern Asia, Europe, and Africa) from 1000 Genome Project were included in the library. 3,074 reference sequences which have a variant for test and randomly selected 2,792 reference sequences with canonical REST motifs were selected and scrambled motif sequences. The 21-bp motifs for these sequences were randomly shuffled and then their binding score of REST checked by using SPRy-SARUS (ver2.0.1) in HOCOMOCO v11 by using a cut-off score of 5. Scrambled sequences which had a REST binding motif detected by SPRy-SARAS starting from 75-110th nucleotides were randomly scrambled again. The random genomic controls from the benchmarking set were also included in the whole genome RE1 library.

### Log additive modeling

Log additive model and its score were calculated by using LinearRegression from Scikit learn (version 0.24.2). The coefficients and fitness scores were resulted as below:

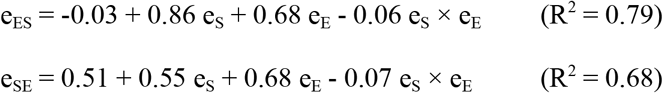

### Target gene expression

Target genes of RE1 were selected as the nearest gene by using the closest-features function of BEDOPS (version 2.4.40) ^58^. RNAseq of K562 from ENCODE (ENCFF088RDE) and Ensembl gene annotations (GRCh37.87) were used. Genes with no expression were removed before analysis.

### ChIP-seq dataset

All analysis with ChIP-seq data was done with the GRCh38 human genome. Reference sequences in the tested library were converted to GRCh38 using liftOver (ENCODE). Accession numbers of ChIP-seq data in ENCODE database are listed in Supplemental Table 11. Overlap between tested elements and ChIP-seq peaks is detected by bedtools (options: -F 0.5 -f 0.5 -e). Heatmap is plotted by using deepTools ^59^ (version 3.5.1, options: --colorMap RdBu --whatToShow ‘heatmap and colorbar’ --zMin −4 --zMax 4 -heatmapWidth 10) after making matrix (options: --referencePoint center -b 1000 -a 1000 --skipZeros -p 4) from bigwig files with accession number ENCFF407OAJ, ENCFF090ZAX, ENCFF313DTO, ENCFF857APX and ENCFF065PDS.

### Identifying half sites and non-canonical motif

Half sites were identified using FIMO. The 1st-9th and 12th-21st nucleotides of REST binding motif (JASPAR MA0138.2) were used for left and right half sites respectively. REST binding sites with two half sites separated with a spacer under 100 bp were categorized according to orientations of the two half sites.

### Piecewise regression

For piecewise regression between predicted binding score and log skew for reference and scrambled, segmented ^60^ was used (ver. 1.3-4, psi=c(20)). The results of the regression and R^2^ are in Supplemental Table 10. The average score of junctions, 20.86, was calculated without the two outliers (En02 and En11 for GM12878) and used to separate weak and strong motifs.

### Genome wide variant overlapping to RE1

All single-nucleotide substitutions from gnomAD v3.1.1 were checked to determine whether both alleles overlap with a REST binding motif (MA0138.2) by using FIMO with a ±20 bp window with a threshold score of 20.89 in either allele. Frequency of variants were called using MafDb.gnomAD.r3.0.GRCh38. Random variants were generated using 5 runs of Mutation-simulator ^61^ (Ver 2.0.3, options: -sn 0.001) followed by the collection of the percentage of nucleotide substitutions by referring to the proportion of all gnomAD variants from chromosome 1.

## Data and code availability

Datasets supporting this manuscript are available at NCBI GEO (Accession ID: GSE196171). Code supporting this manuscript is available on GitHub (general processing pipeline for MPRAduo: https://github.com/tewhey-lab/MPRAduo, data analysis: https://github.com/tewhey-lab/duoREST).

## Acknowledgement

We gratefully acknowledge the contribution of Ryan Lynch and Genome Technologies Service at The Jackson Laboratory. This work was funded by grants R00HG008179 and R35HG011329 awarded to R.T..

## Author contribution

K.M. and R.T. conceived the study. K.M. performed MPRA with the help of D.B. and S.K. K.M., H.B.D. and R.C performed data analysis. K.M., H.B.D., R.C., and R.T. wrote the manuscript. All authors have read and approved the manuscript.

## Supplemental Tables

Supplemental Table 1: Primers for library construction and sequencing

Supplemental Table 2: Sequences tested in this study

Supplemental Table 3: Summary of enhancers

Supplemental Table 4: Results of benchmarking libraries for run1 and run2

Supplemental Table 5: Summary of TF binding groups in benchmarking libraries

Supplemental Table 6: Results of whole genome RE1 library

Supplemental Table 7: List of ChIP-seq dataset from ENCODE

Supplemental Table 8: Predicted binding scores of tested motifs

Supplemental Table 9: Results of motif contribution

Supplemental Table 10: Summary of piecewise linear regression

Supplemental Table 11: Summary of colocalization of TF and accession numbers of ChIP-seq dataset

Supplemental Table 12: Results of allelic skew

