## Supplementary Figures for "Whole genome functional characterization of RE1 silencers using a modified massively parallel reporter assay"

### Supplemental Figure 1

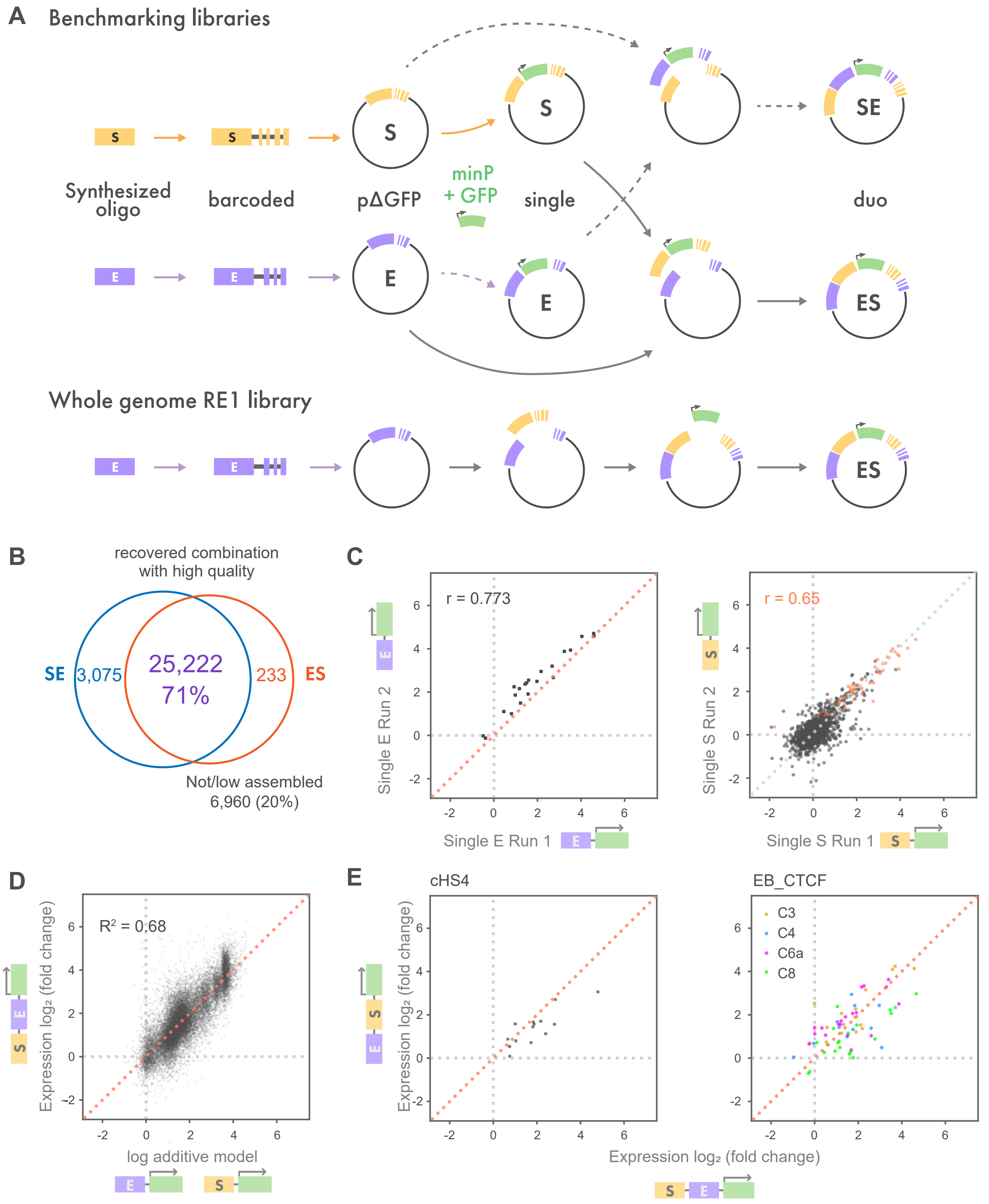


(A) Detailed workflow of MPRAduo construction. For the benchmarking libraries, oligos are synthesized, barcoded, and cloned as single libraries (pΔGFP), then GFP ORF and a minimal promoter (minP) are inserted between test element and barcode. Next, the oligo and barcode sequences including GFP ORF and minP cassette are amplified and cloned into a reciprocal pΔGFP library. For the whole genome RE1 library, pΔGFP for E library is cloned and then S-oligo cassettes with barcodes are inserted. Next, GFP ORF and a minimal promoter (minP) are inserted between S elements and barcodes. (B) Overlap of the combinations of S and E detected in the benchmarking duo libraries with high quality read counts. (C) Correlation of normalized expression level (log2 of the mRNA/plasmid DNA ratio) between two runs of single libraries. r indicates Pearson’ s correlation of active elements. (D) Correlation between log additive model of single libraries and their observed duo values in ES library. (E) Correlation of normalized expression level of cHS4 and CTCF binding sites in known EB in duo libraries.

### Supplemental Figure 2

**A**

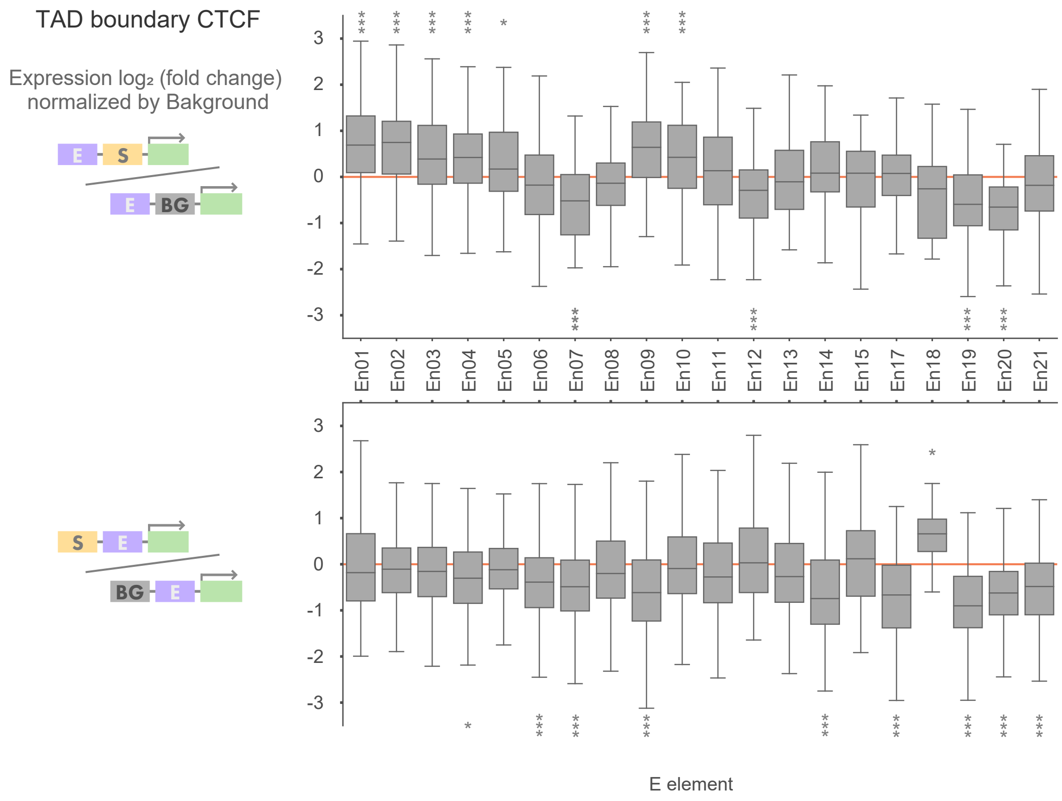


**B**

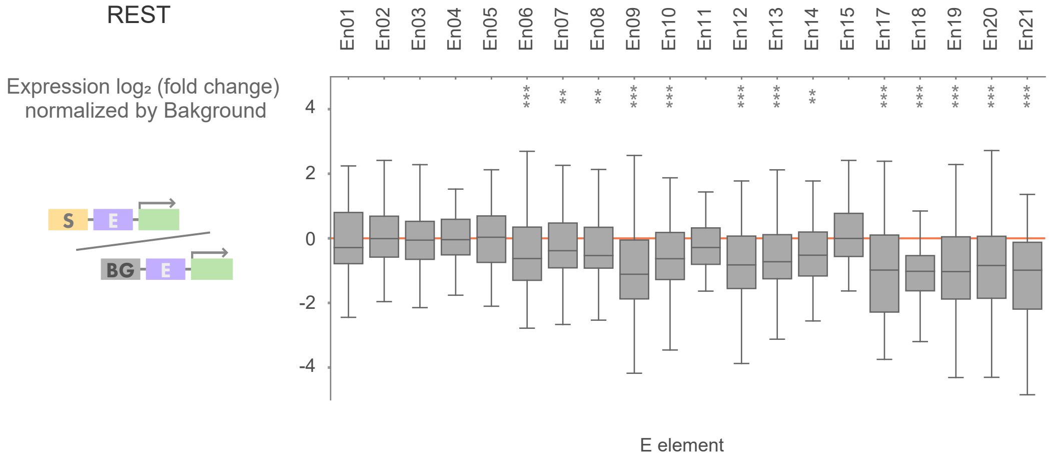


MPRA activity of the combinations of CTCF at TAD boundary (A) or RE1 (B) and enhancers normalized by the distribution of the corresponding combinations of RE1 and the random genomic control. *: adjp<0.05, **: adjp<0.01, ***: adjp<0.001 by U-test compared with background controls corrected using BH.

### Supplemental Figure 3

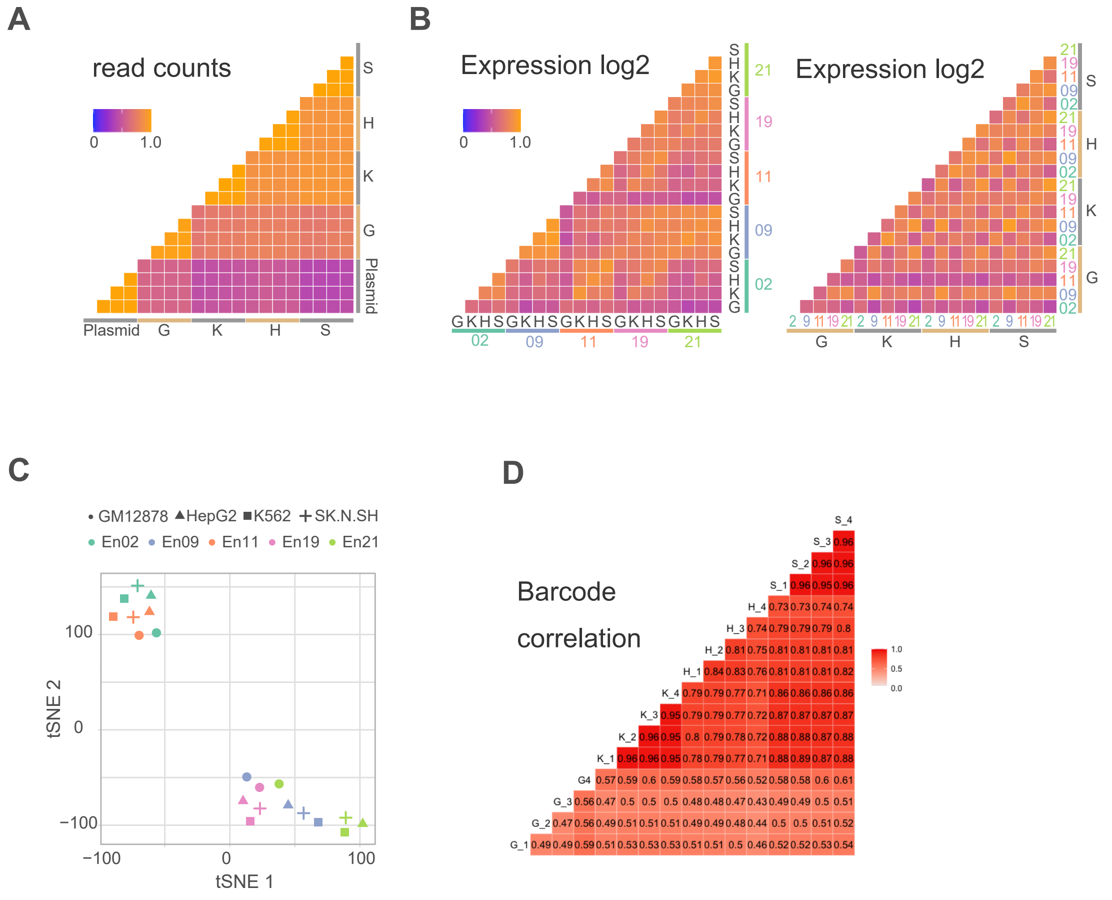


(A) Correlations of read counts between replicates of mRNA and plasmid DNA counts. The color bar indicates Pearson' s correlation. (B) Correlations of expression level (log2 of the mRNA/plasmid DNA ratio) between combinations of cell and enhancer. Same results are sorted by enhancer (left) or cell type (right). (C) tSNE plot by using expression levels of the RE1 with canonical motif. 20 data sets are normalized. (D) Correlations of barcode variations between replicates of mRNA.G: GM12878, K: K562, H: HepG2, S: SK-N-SH

### Supplemental Figure 4

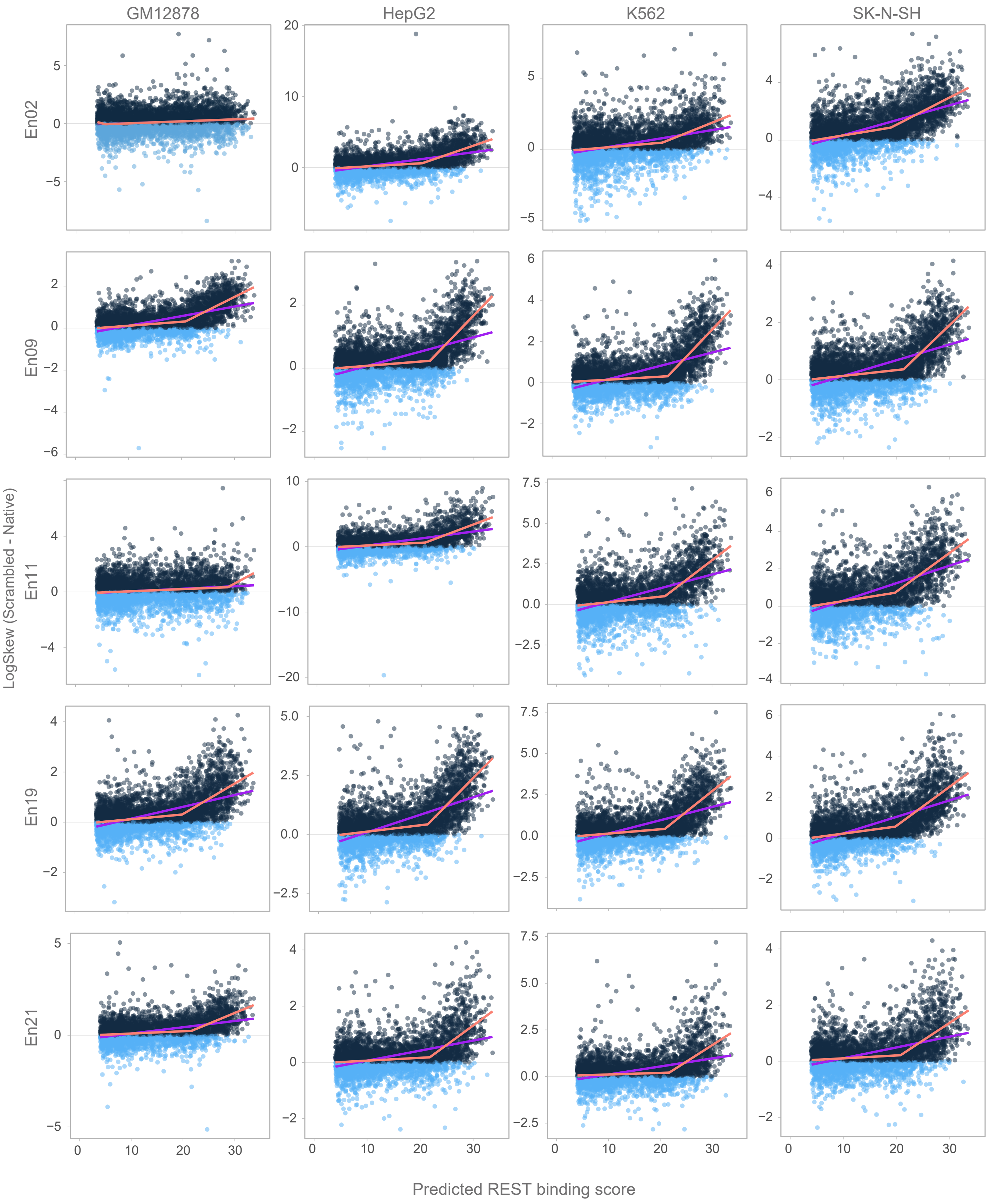


Correlation between predicted binding score of REST and motif contribution (log2(Scrambled / Native)). The purple lines indicate linear regression and the orange lines indicate piecewise linear regression.

### Supplemental Figure 5

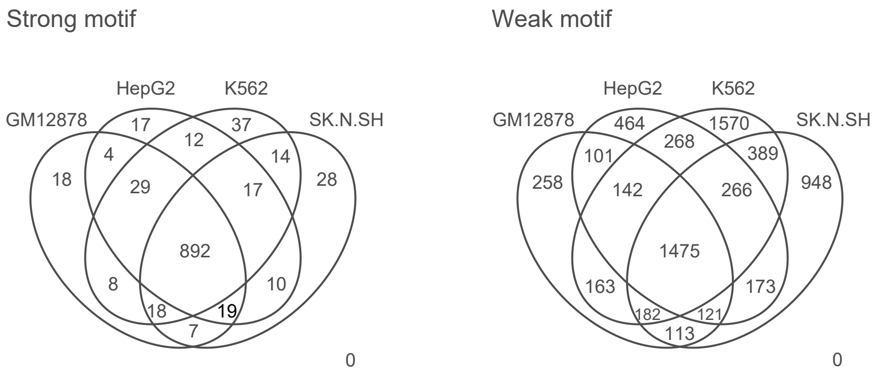


Venn diagram of colocalization of the ChIP-seq peaks in the 4 cell lines for tested RE1s with strong or weak motif.

### Supplemental Figure 6

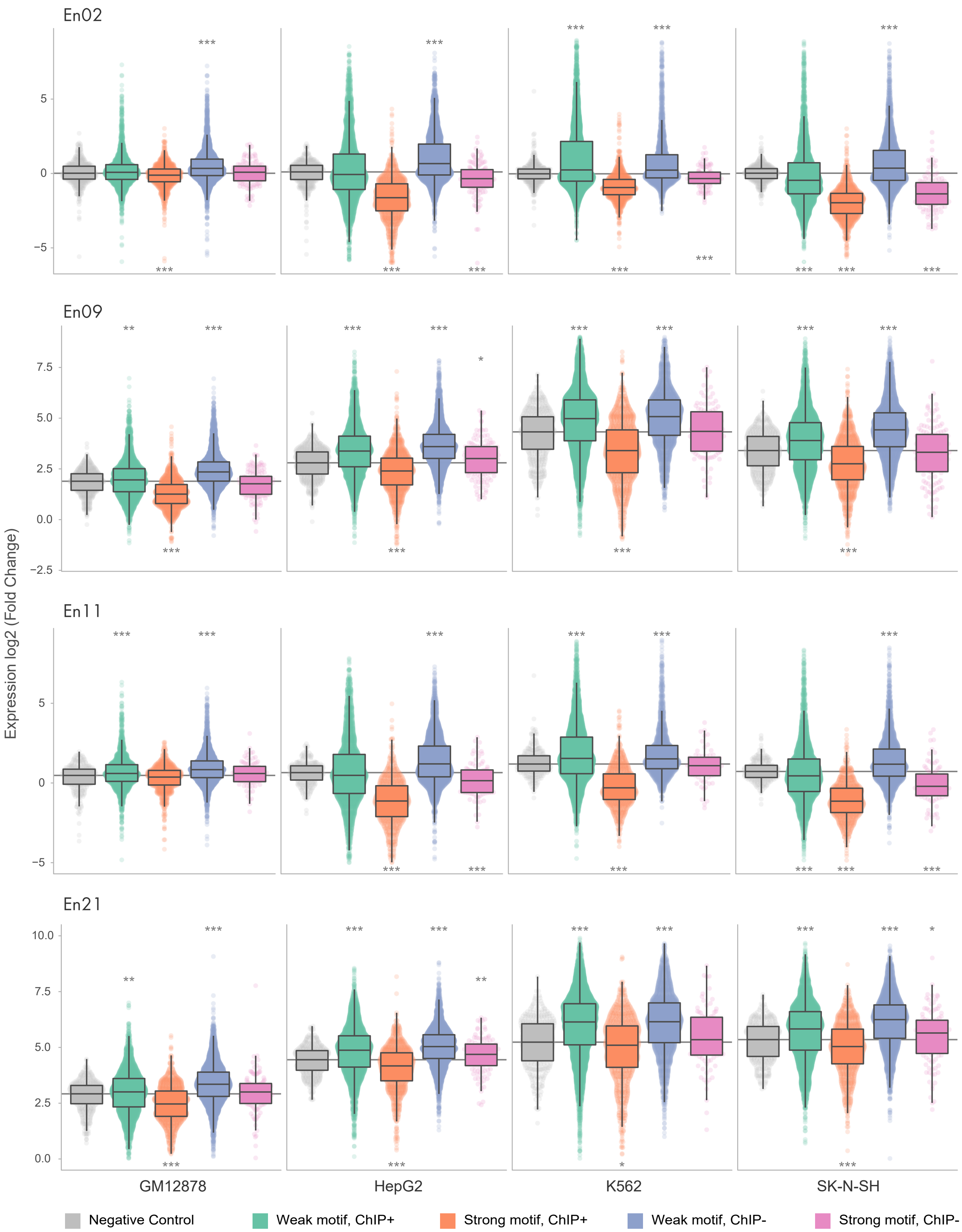


MPRAduo activity for each E element-cell type combination. Specific-ON indicates RE1s bound by REST in each cell type and Specific-OFF indicates RE1s not bound by REST in the observing cell type but bound in other cell type(s). Grey line indicates the median of the negative control. *: adjp<0.05, **: adjp<0.01, ***: adjp<0.001 by U-test compared with negative controls corrected using BH.

### Supplemental Figure 7

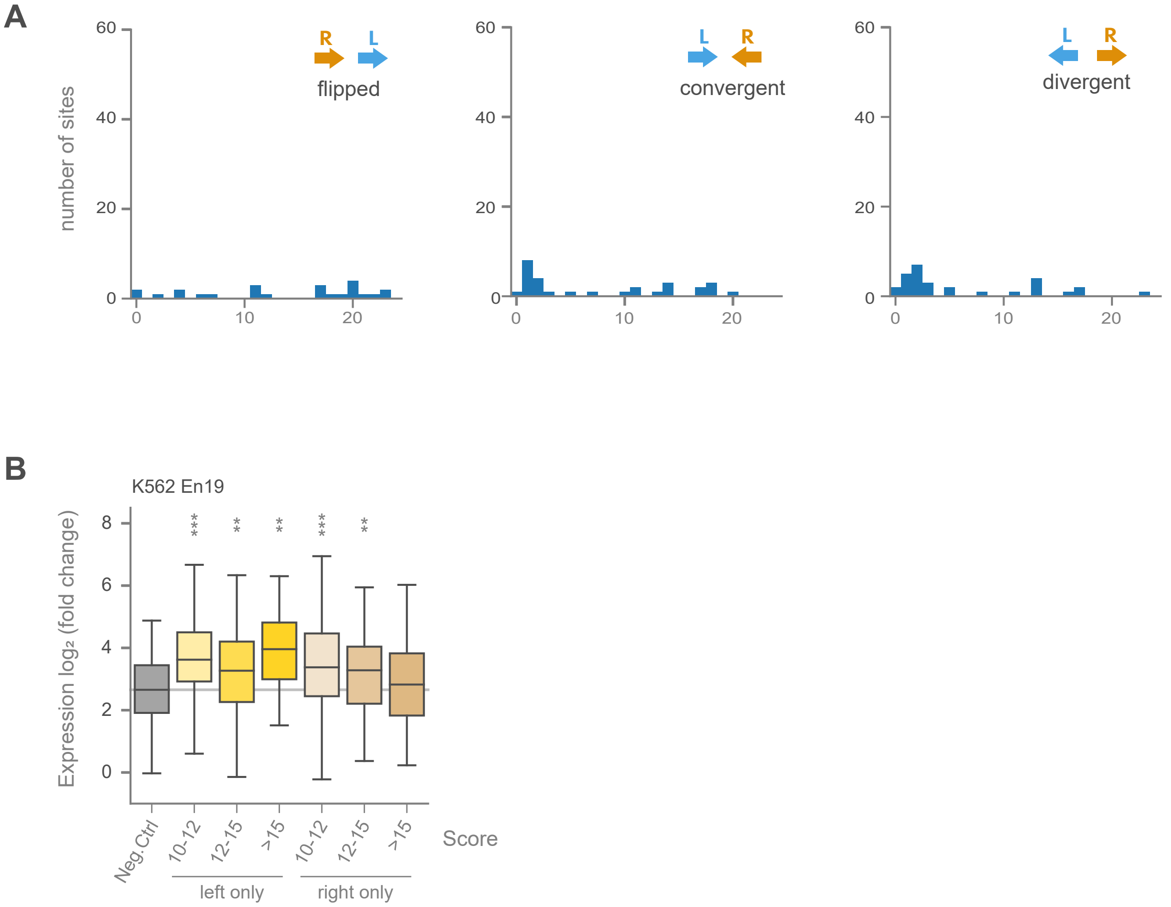


(A) Distribution of spacer length between two half sites aligned in flipped, convergent or divergent manner in the tested library. (B) MPRAduo activity of half binding sites with different binding scores in K562 with En19. Grey line indicates the median of the negative control **: adjp<0.01, ***: adjp<0.001 by U-test compared with negative controls corrected using BH.

Supplementary Figure 8

(A) Categorizing plot using TF ChIP-seq in HepG2 and reporter activity in HepG2 with En19. Red dots are TFs with significant differences despite REST colocalization. Blue and green dots are TFs with significant differences only with or without REST respectively (p<0.05 by U-test). (B) Enrichment of TF localization in HepG2 at RE1 with strong motifs compared to all tested canonical RE1. Red dots are significantly enriched TFs (FDR < 0.05 by Fisher' s exact test). (C) Correlation between number of localized category (i) TFs and expression level of MPRAduo with En19 in K562. (D) Correlation between number of localized category (i) TF and REST motif contribution with En19 in K562 (box plot, left axis). Red line indicates the proportion of REST binding sites of each bin in the genomic context of K562 (right axis). (E) Number of category (i) and category (ii) TFs binding in the genomic context of HepG2 and its effect on expression level of MPRAduo with En19 in HepG2. (F) Number of category (i) and category (ii) TFs binding in the genomic context of HepG2 and REST motif contribution (box plot, left axis) measured by MPRAduo with En19 in HepG2. Red line indicates the proportion of REST binding sites of each bin in the genomic context of HepG2. (G) Binding property of REST and cofactors in the genomic context of HepG2 and its effect on expression level of MPRAduo with En19 in HepG2. (H) Binding property of REST and cofactors in the genomic context of HepG2 and its effect on motif contribution of MPRAduo with En19 in HepG2. *: p<0.05, **: p<0.01, ***: p<0.001 by U-test compared with negative controls or RE1s bound by one TF corrected using BH.

### Supplemental Figure 8

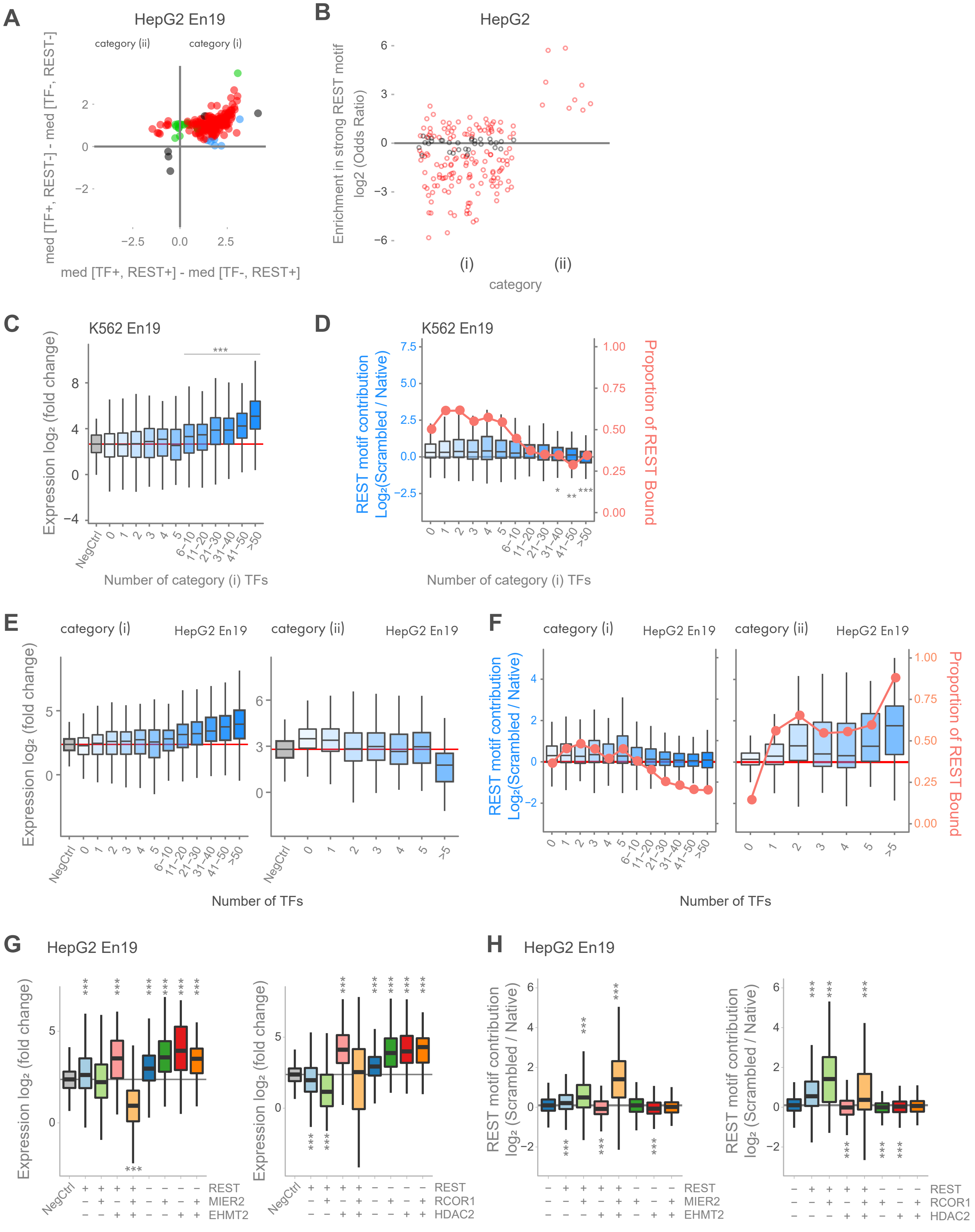

### Supplemental Figure 9

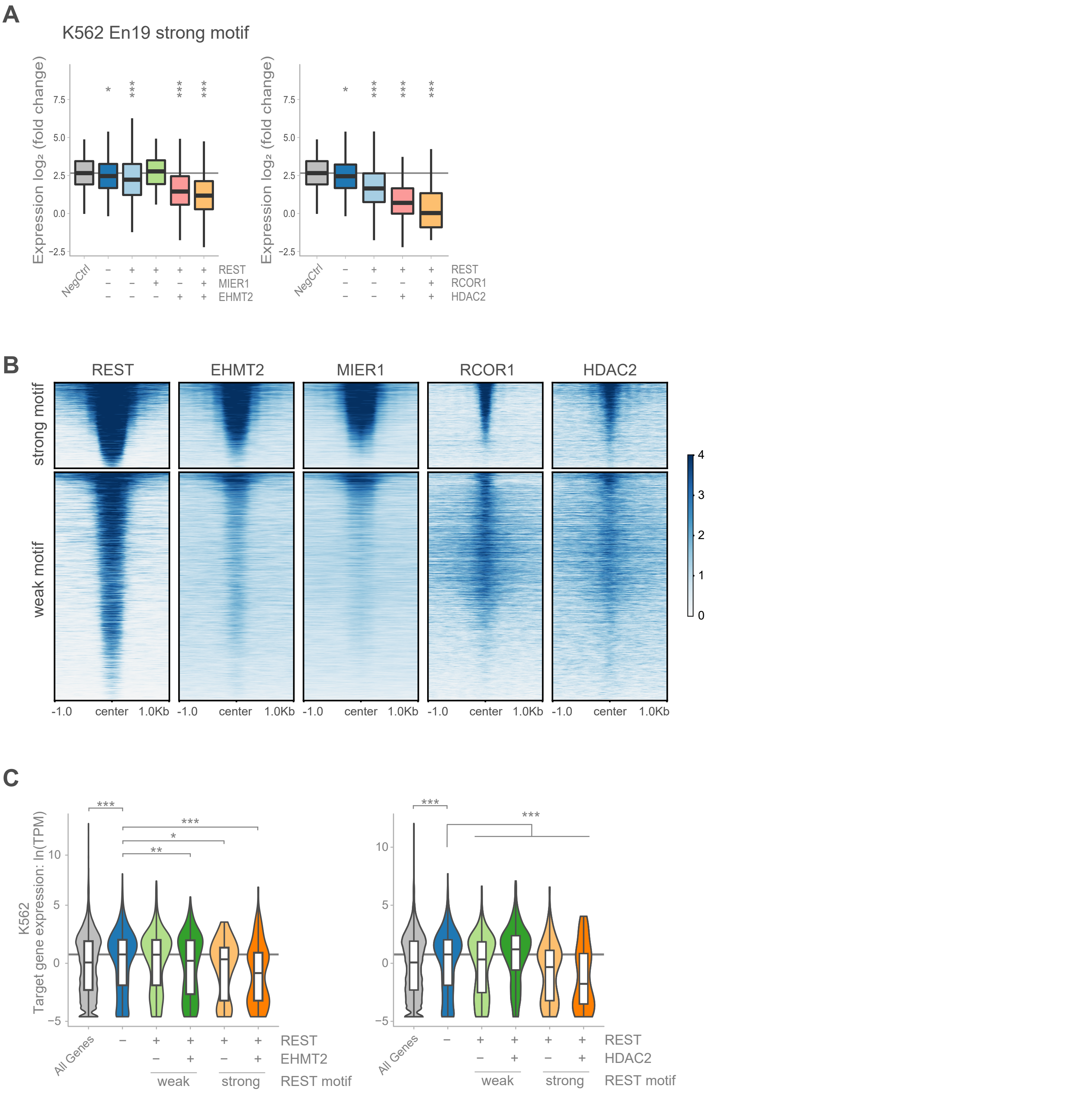


(A) Binding property of REST and cofactors in the genomic context of K562 on RE1 with strong motif and its effect on repressive activity of MPRAduo with En19 in K562. (B) Heatmap of ChIP-seq peaks for K562 from - 1 kb to + 1 kb of the center of tested RE1s with strong or weak motifs. (C) Effect of motif strength and cofactor binding on expression level of their target gene in K562. *: p<0.05, **: p<0.01, ***: p<0.001 by U-test compared with negative controls or genes targeted by specific-OFF RE1s corrected using BH.

### Supplemental Figure 10

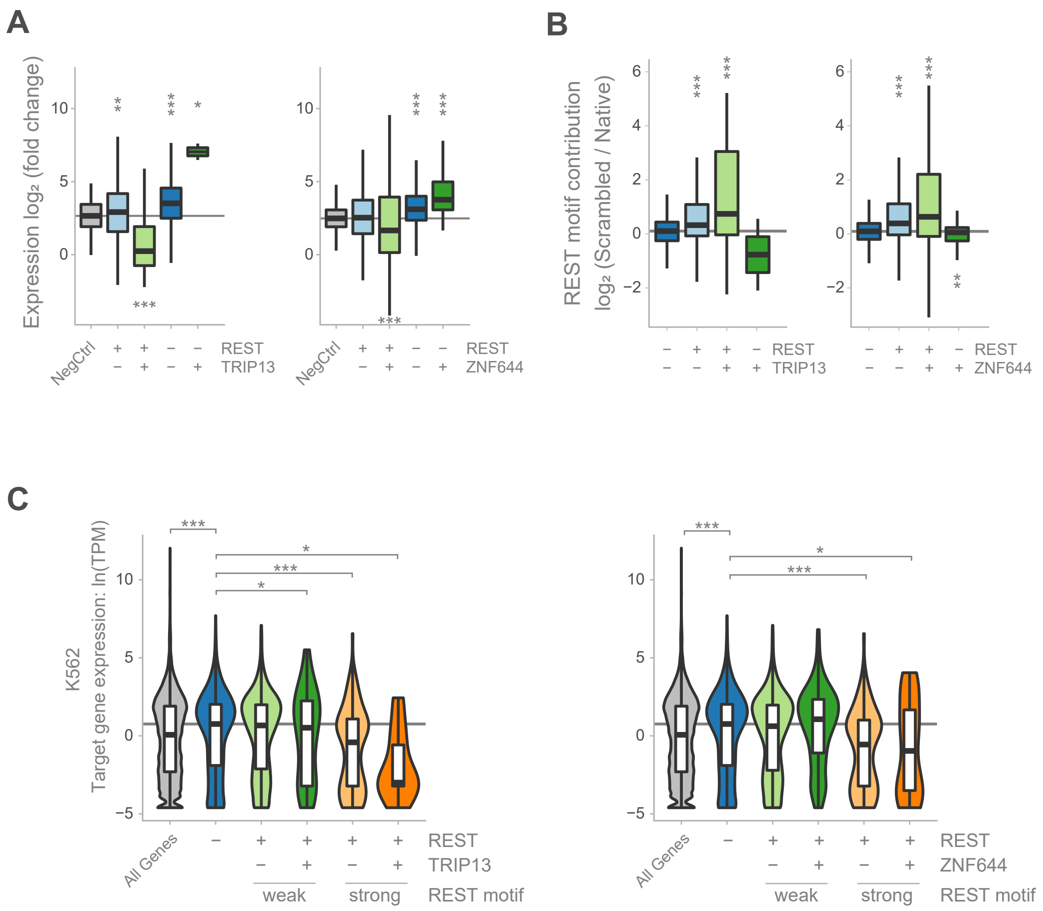


(A) Binding property of TRIP13 or ZNF644 with REST in the genomic context of K562 and its effect on expression level of MPRAduo with En19 in K562. (B) Binding property of TRIP13 or ZNF644 with REST in the genomic context of K562 and its effect on REST motif contribution with En19 in K562. (C) Effect of REST motif strength and TRIP13 or ZNF644 binding on expression level of their target gene in K562 shown in log of TPM (Transcription per kilobase million). *: p<0.05, **: p<0.01, ***: p<0.001 by U-test compared with negative controls (A), double negative group (B) or genes targeted by specific-OFF RE1s (C) corrected using BH.

### Supplemental Figure 11

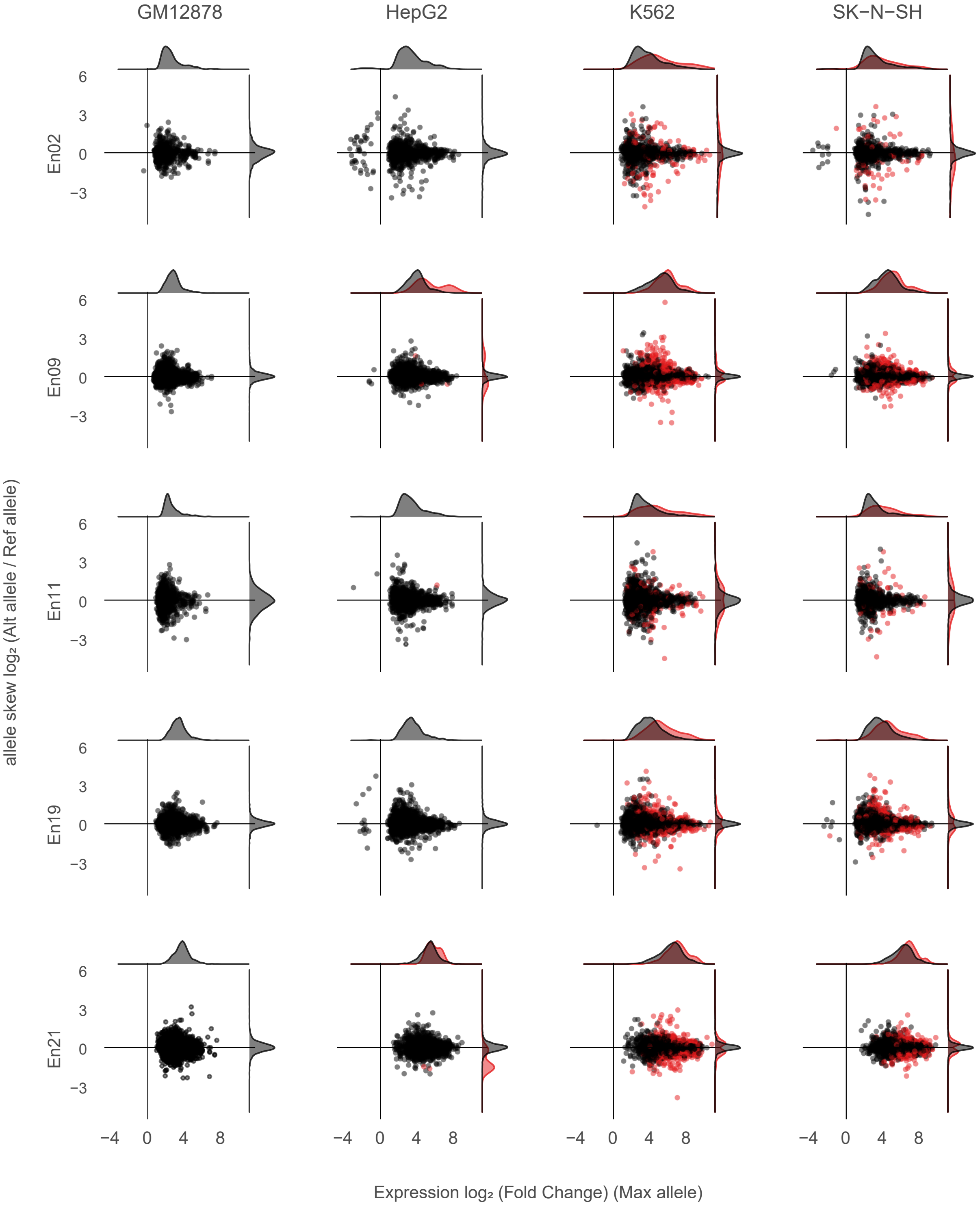


Distributions of expression level (x-axis) of maximum allele and allelic skew (y-axis). Red dots are emVar with FDR < 0.05.

### Supplemental Figure 12

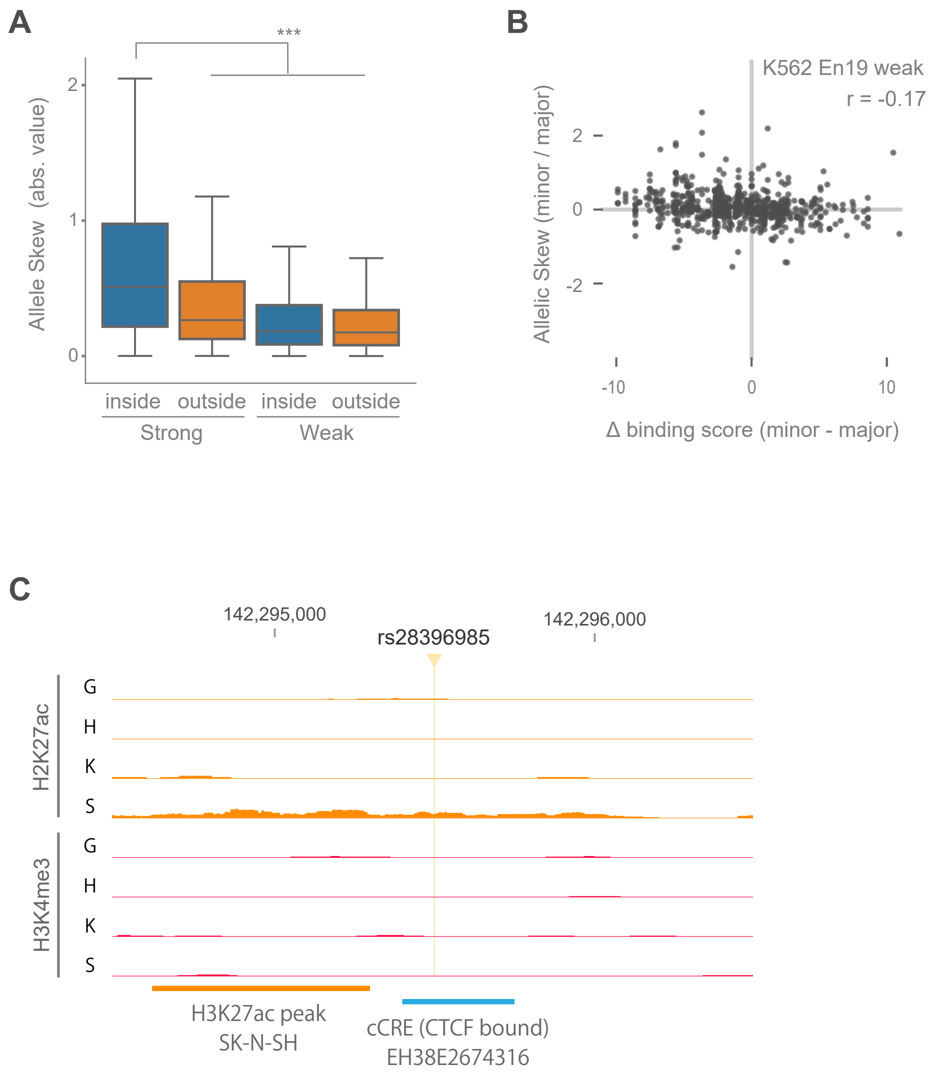


(A) Comparing absolute value of allelic skew between strong/weak motifs and inside/outside of the motif. ***: $p<0.001$ by U-test (B) Correlation of allelic skew and difference of binding scores of allele at weak motifs. allelic skew with En19 in K562 is plotted against the difference of predicted binding score between alleles. (C) Position of rs28396985 based on GRCh38, ChIP-seq signals of H3K27ac and H3K4me3, called peak of H3K27ac and CRE annotation by SCREEN in TSNARE1 locus. G: GM12878, H: HepG2, K: K562, S: SK-N-SH. The y-axis is from 0 to 32 for all tracks.

### Supplemental Figure 13

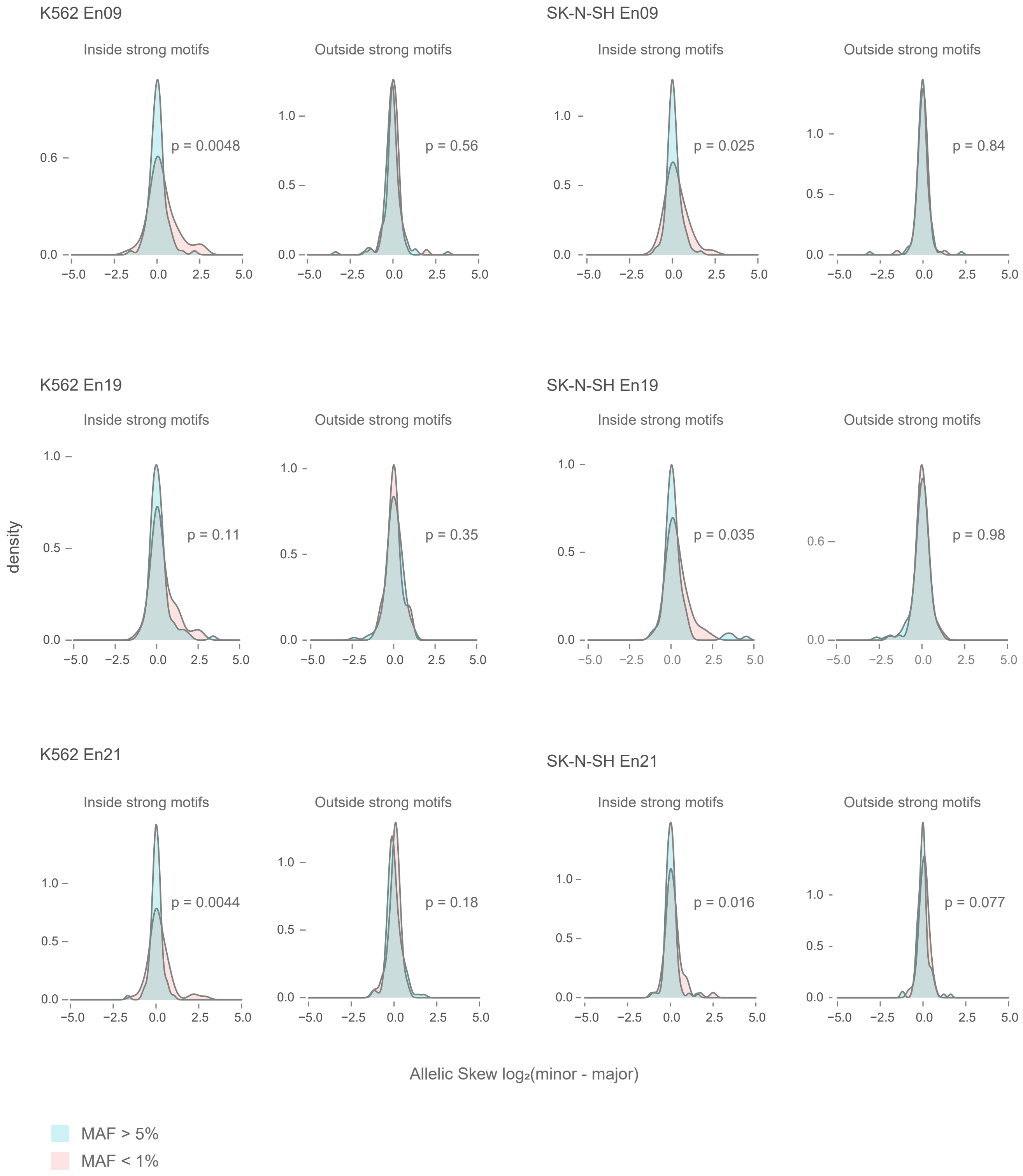


Distribution of allelic skew in different MAF. p-values are calculated by U-test.

#### Supplemental Fig. 14

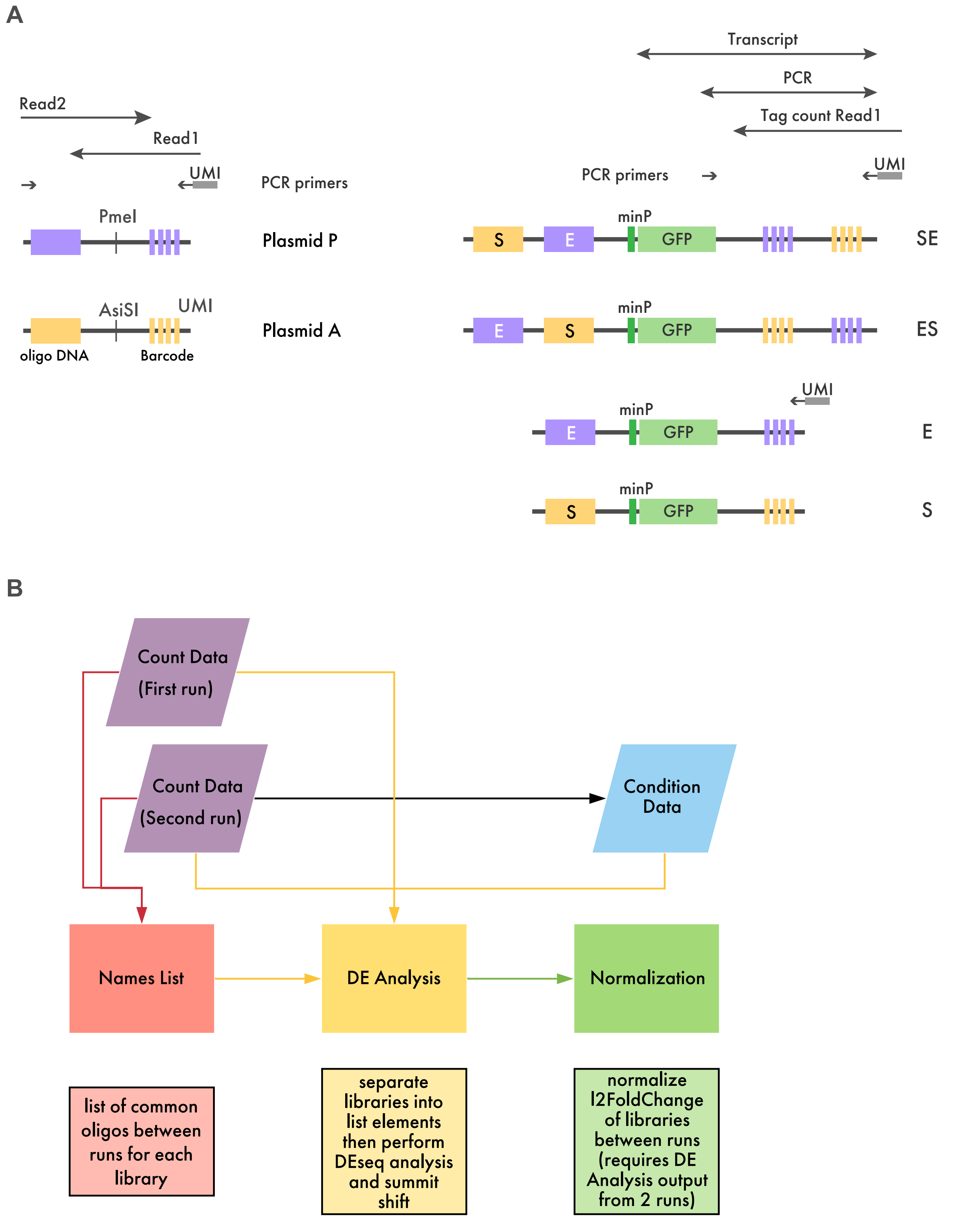


(A) Schematic structure of PCR product, transcript, and sequencing reads of single and duo libraries. (B) Workflow of read count analysis including pre-process and full analysis.
